# An interplay of multilayered regulatory mechanisms underlies the RNA solubility phase transition during vertebrate oocyte-to-embryo transition

**DOI:** 10.64898/2026.11.11.765953

**Authors:** Sijie Chen, Meng Ma, Danping Chen, Minghui Lu, Zhou Wan, Hyojeong Hwang, Samuel Hatlestad, Seunghee Choi, Harsh Patel, Kavya Panjrath, Aidan Adachi, Kun Wang, Wenyan Mei, Boxuan Zhao, Jing Yang

**Affiliations:** Department of Comparative Biosciences, University of Illinois at Urbana-Champaign, Urbana, IL 61802, USA; Department of Cell and Developmental Biology, University of Illinois at Urbana-Champaign, Urbana, IL 61801, USA; Department of Biochemistry and Biophysics, Perelman School of Medicine, University of Pennsylvania, Philadelphia, PA 19104, USA; School of Molecular and Cellular Biology, University of Illinois at Urbana-Champaign, Urbana, IL 61801, USA

**Keywords:** RNA phase transition, RNA insolubility, poly(A) tail, m^6^A, oocyte-to-embryo transition, zebrafish, *Xenopus*

## Abstract

Dynamic RNA-protein interactions influence RNA transport, storage, stability, and activity, and are fundamental to a broad range of biological processes. During the oocyte-to-embryo transition (OET), fully differentiated gametes fuse and regain totipotency. Throughout the OET, the transcription remains silent. Spatiotemporal regulation of gene expression relies on the dynamic interactions between maternal transcripts and various protein complexes. Despite its fundamental importance, RNA-protein interaction dynamics at the global level during the OET remain poorly understood. Here, we leverage fractionation RNA-seq to profile RNA insolubility, a proxy for the interaction between RNAs and protein complexes, during zebrafish OET. We identify transcripts that transition between the soluble (relatively protein-free) and insoluble (heavily protein-associated) states and demonstrate that these phase transitions are driven by ribosome-dependent and ribosome-independent mechanisms. We demonstrate that the capacity of maternal RNAs to engage with protein complexes changes dynamically during the OET through remodeling of *cis*-regulatory features. We developed a machine-learning framework and uncovered multiple *cis*- and *trans*-regulatory features that may contribute to RNA insolubility. To validate these predictions, we examined poly(A) tail length and m^6^A dynamics. We found that poly(A) tail length is associated with both ribosome-dependent and ribosome-independent RNA phase transitions. In parallel, fractionation m^6^A-pulldown RNA-seq revealed highly dynamic changes in the m^6^A landscape during the OET. A correlation between m^6^A modifications and RNA insolubility is gradually established during the OET. Together, our findings uncover a multilayered regulatory mechanism governing RNA phase transitions during the OET, providing fundamental insights into post-transcriptional regulation of gene expression during development.

**Article Summary:** The spatiotemporal regulation of gene expression during the oocyte-to-embryo transition (OET) relies on dynamic interactions between maternal transcripts and various protein complexes, yet RNA-protein interaction dynamics at the global level during the OET remain poorly understood. By analyzing RNA insolubility, a proxy for RNA-protein interactions, we identify transcripts that transition between the soluble (relatively protein-free) and insoluble (heavily protein-associated) states. We demonstrate that RNA phase transitions during the OET are regulated by an intricate interplay among static sequence-specific *cis*-elements, dynamic RNA features, and *trans*-acting factors. Our work has provided fundamental insights into post-transcriptional regulation of gene expression during development.

## Introduction

Post-transcriptional mechanisms play pivotal roles in the precise regulation of gene expression across a wide range of biological processes. These mechanisms rely on dynamic interactions between RNAs and RNA-binding proteins (RBPs), which govern RNA maturation, stability, subcellular localization, and activity. RNA-protein interactions can be regulated in *trans* by controlling the expression and post-translational modifications of RBPs. They can also be influenced by various *cis*-regulatory mechanisms, including sequence-specific *cis*-elements in the RNA, poly(A) tail dynamics, and epitranscriptomic modifications. Despite their importance, how dynamic RNA-protein interactions are precisely coordinated across diverse biological processes remains largely unclear.

The oocyte-to-embryo transition (OET) is a unique developmental process in which fully differentiated gametes fuse and regain totipotency. During the OET, oocyte maturation, ovulation, fertilization, and zygotic genome activation (ZGA) occur sequentially, transforming the dormant oocyte into a rapidly developing embryo. This transition is primarily regulated by maternal factors deposited in the oocyte during oogenesis and proceeds in the absence of transcription (Conti and franciosi 2018; Schultz *et al*. 2018). It is well known that although oocytes store a large number of maternal transcripts, only specific subsets are translated at defined times and locations to coordinate developmental events during the OET (Wang *et al*. 2010; Winata and korzh 2018; Yang *et al*. 2020; Hu *et al*. 2022; Xiong *et al*. 2022; Zhu *et al*. 2022; Iyyappan *et al*. 2023; Zhang *et al*. 2023; Conti and kunitomi 2024). The mechanisms by which the translational machinery selectively activates a subset of mRNAs while maintaining the majority of the maternal transcriptome in a silent state during the OET are poorly understood.

Recent studies have uncovered several regulatory mechanisms that are fundamental to gene expression during the OET (Conti and franciosi 2018; Winata and korzh 2018; Sha *et al*. 2019; Ermisch and wood 2024; Lorenzo-orts and pauli 2024). In some vertebrate species, maternal RNAs are stored on the surface of cellular organelles (Cheng *et al*. 2022; Hwang *et al*. 2022) and are maintained in a dormant state in fully grown oocytes (Ermisch and wood 2024; Lorenzo-orts and pauli 2024). Upon oocyte maturation, these organelle-associated dormant RNAs are released into the cytoplasm (Cheng *et al*. 2022; Hwang *et al*. 2022), where they gain access to the translational machinery or other protein complexes that fine-tune gene expression. Unlike in other biological contexts, however, the translational machinery itself is constrained during the OET. In many species, the levels of ribosomes and poly(A)-binding proteins (PABPs), key components of the translation machinery, are tightly regulated. Previous studies have shown that ribosome biogenesis is suppressed during the OET and resumes only after the ZGA (Piko and clegg 1982; Gao *et al*. 2017; Hu *et al*. 2022; Xiong *et al*. 2022; Zhang *et al*. 2022a; Zhu *et al*. 2022; Zhang *et al*. 2023; Galatidou *et al*. 2024). Furthermore, approximately 70-80% of ribosomes are inactive in growing mouse oocytes and ovulated eggs (Bachvarova and de leon 1977; Bachvarova *et al*. 1981; De leon *et al*. 1983). In zebrafish and *Xenopus,* maternal ribosomes are simultaneously bound by inhibitory factors, including eIF5a, eEF2b, Habp4, and Dap1, leading to ribosome dormancy (Leesch *et al*. 2023). Moreover, ePAB, the PABP predominantly expressed during the OET (Voeltz *et al*. 2001; Seli *et al*. 2005; Esencan *et al*. 2019; Ozturk 2019), is expressed at levels that are limiting relative to the number of available poly(A) sites (Xiang and bartel 2021). Consequently, maternal transcripts must compete for the limited pool of translational machinery throughout the OET. Given that tens of thousands of maternal RNAs are crowded in the oocyte, precisely regulated RNA-protein interactions are thus essential for orchestrating temporospatial gene expression during the OET.

RNA-protein interactions are controlled by an interplay of regulatory mechanisms functioning at both the *cis* and *trans* levels. Many RBPs are dynamically expressed or post-translationally modified during the OET (Conti and franciosi 2018; Winata and korzh 2018; Thomalla and wolfner 2025). Meanwhile, although maternal RNAs contain many static sequence-specific *cis*-elements, tail dynamics mediated by the addition or removal of non-templated nucleotides (Yu and kim 2020; Liu and lu 2024), epitranscriptomic mechanisms, i.e., RNA chemical modifications (Nachtergaele and he 2018), and changes in RNA secondary structure during the OET all heavily influence the ability of maternal RNAs to engage *trans*-acting factors. It is well-known that the poly(A) tail length of maternal RNAs is dynamically regulated by cytoplasmic polyadenylase and deadenylases during the OET (Barckmann and simonelig 2013). As the interaction between PABP and poly(A) tail facilitates translation, the length of poly(A) tails of maternal RNAs correlated well with their ribosome occupancy (Subtelny *et al*. 2014; Xiang and bartel 2021). It has also been reported that uridylation of mRNA tails, catalyzed by terminal uridylyltransferases TUT4/7, plays a crucial role in controlling the stability of maternal RNAs (Chang *et al*. 2018). Epitranscriptomic mechanisms have been extensively studied in the context of the OET as well. These include m^6^A modifications (Zhao *et al*. 2017; Zhang *et al*. 2022b; Wang *et al*. 2023; Li *et al*. 2025; Han *et al*. 2026; Iyyappan *et al*. 2026; Su *et al*. 2026), which regulate the decay and translation of maternal RNAs during the OET, and m^5^C (Yang *et al*. 2019; Liu *et al*. 2022), which regulates the stability of maternal RNAs. Intriguingly, not only are these modifications highly dynamic during the OET (Liu *et al*. 2022; Li *et al*. 2025; Han *et al*. 2026; Iyyappan *et al*. 2026; Su *et al*. 2026), but some of their reader proteins are also dynamically expressed (Xiao *et al*. 2023), providing pivotal mechanisms for dynamic RNA-protein interactions. Nevertheless, owing to the lack of appropriate high-throughput technologies, transcriptome-wide analyses of RNA-protein interaction dynamics during the OET have remained largely unexplored.

We recently developed a fractionation RNA-seq protocol suitable for profiling RNA-protein interaction dynamics during the OET (Hwang *et al*. 2022; Hwang *et al*. 2023; Hwang *et al*. 2024). This method exploits the principle that RNAs associated with large protein complexes exhibit higher molecular weights and therefore sediment into the pellet after centrifugation, whereas relatively free RNAs remain in the supernatant. Quantifying the distribution of transcripts between the soluble (supernatant) and insoluble (pellet) phases thus provides an efficient means of quantitatively assessing RNA-protein interactions. By investigating the RNA insolubility, i.e., the proportion of a transcript distributed in the pellet after centrifugation, we recently reported that maternal transcripts transition between the soluble and insoluble phases during *Xenopus* OET. In particular, the insoluble-to-soluble (I-S) phase transition of germline RNAs, mediated by their dissociation from germ plasm, regulates the stability of germline RNAs (Hwang *et al*. 2023). Similar fractionation-based approaches have been applied to investigate the stress response in budding yeast (Glauninger *et al*. 2025) and phosphorylation-dependent RNA-protein interactions in mammalian cells (Sridharan *et al*. 2022).

In the present study, we characterized RNA-protein interaction dynamics during zebrafish OET. In zebrafish, zygotically expressed microRNAs play a critical role in the clearance of maternal RNAs (Giraldez *et al*. 2006). Because zygotic transcription is absent and RNA turnover remains limited prior to ZGA, the level of most maternal RNAs is largely stable during zebrafish OET (Bhat *et al*. 2023), providing a powerful model system for dissecting RNA-protein interaction dynamics. We employed fractionation RNA-seq to systematically profile dynamic RNA-protein interactions during zebrafish OET. Our results reveal that an intricate interplay among static sequence-specific *cis*-elements, dynamic RNA features such as poly(A) tail remodeling and epitranscriptomic modifications, and *trans*-acting factors collectively determines RNA-protein interaction dynamics and fine-tunes gene expression during the OET.

## Materials and methods

### Danio rerio

Zebrafish procedures were approved by the University of Illinois at Urbana-Champaign Institutional Animal Care and Use Committee (IACUC) under animal protocol #25153 and performed in accordance with the recommendations of the Guide for the Care and Use of Laboratory Animals of the National Institutes of Health. The female Tübingen strain was purchased from ZIRC (Zebrafish International Resource Center). Embryos at the 2-, 32-, and 256-cell stage were collected from a mating cross of wild-type zebrafish. After embryo collection, the female fish was given one week for recovery and used again to collect unactivated eggs. One more week later, the same female fish was used for oocyte collection as described (Lee and marlow 2019). Briefly, ovarian tissues were dissected and dissociated by 15 mg/ml collagenase treatment for 30 min at room temperature (RT). Fully-grown oocytes were collected manually.

### Xenopus laevis

*Xenopus* procedures were approved by the University of Illinois at Urbana-Champaign Institutional Animal Care and Use Committee (IACUC) under animal protocol #26082 and performed in accordance with the recommendations of the Guide for the Care and Use of Laboratory Animals of the National Institutes of Health. Oocytes were collected from ovarian tissues by manual defolliculation and cultured in the oocyte culture medium (OCM) (Houston 2018). In order to induce oocyte meiotic maturation, stage VI oocytes were cultured in the OCM containing 2 µM progesterone and incubated at 18 °C overnight. Oocytes and mature eggs were collected for further analysis. For microinjection, 16 ng of zebrafish total RNA was injected into *Xenopus* oocytes using a Narishige IM300 microinjector. Injected oocytes were either cultured in the OCM or treated with progesterone.

#### Fractionation during zebrafish OET

The fractionation RNA-seq during zebrafish OET was performed with modifications based on the previously described *Xenopus* fractionation RNA-seq protocol (Hwang *et al*. 2022; Hwang *et al*. 2023; Hwang *et al*. 2024). Briefly, fully-grown zebrafish oocytes were isolated according to a published protocol (Lee and marlow 2019). For fractionation of zebrafish samples, twenty fully-grown oocytes, ovulated eggs, or embryos at the 2-cell, 32-cell or 256-cell stages were homogenized using disposable pestles in 200 µL of ice-cold extraction buffer (150 mM KOAc, 2.5 mM MgOAc2, 20 mM K-HEPES pH 7.4, 0.5% Triton X-100, 2 mM DTT, 1 mM PMSF, 50 µg/mL cycloheximide, and 200 units/mL RNase inhibitor). Lysates were then centrifuged at 10,000 x g for 10 min at 4 °C. After centrifugation, the supernatant was transferred to a new tube as the soluble fractions. The pellet was resuspended in 200 µL of ice-cold extraction buffer as the insoluble fraction.

For ribosome stripping, twenty zebrafish embryos at the 256-cell stage were homogenized using disposable pestles in 200 µL of ice-cold EDTA-containing extraction buffer (150 mM KOAc, 2.5 mM MgOAc2, 20 mM K-HEPES pH 7.4, 0.5% Triton X-100, 2 mM DTT, 1 mM PMSF, 30 mM EDTA, and 200 units/mL RNase inhibitor). Lysates were then centrifuged at 10,000 x g for 10 min at 4 °C. After centrifugation, the supernatant was transferred to a new tube as the soluble fractions, and the pellet was resuspended in 200 µL of ice-cold extraction buffer as the insoluble fraction.

#### Cross-species test of zebrafish RNA insolubility

Total RNAs were extracted from zebrafish fully-grown oocytes and 256-cell stage embryos using TRIzol reagent according to the manufacturer’s instructions. Briefly, twenty fully-grown oocytes or 256-cell stage were lysed in TRIzol. Total RNAs were dissolved in 10 µL of RNase-free water. Then, 16 ng of the RNA mixture was injected into *Xenopus* oocytes. Injected oocytes were treated with progesterone for oocyte maturation. Four oocytes or mature eggs were homogenized in 160 µL of ice-cold extraction buffer and centrifuged at 10,000 x g for 10 min at 4 °C. The supernatant was collected as the soluble fraction. The remaining pellet was resuspended in 160 µl of ice-cold extraction buffer as the insoluble fraction.

#### Library preparation and RNA sequencing

Before RNA extraction, a mixture of spike-in RNAs was added to TRIzol lysate. RNAs were extracted using TRIzol reagent according to the manufacturer’s instructions. For zebrafish fractionation RNA-seq, ribosomal RNAs were depleted using the FastSelect Fish rRNA Removal Kit (Qiagen). Random-primed sequencing libraries were prepared from fractionated RNA using the KAPA HyperPrep Stranded RNA-Seq Library Preparation Kit (Roche). Libraries were then sequenced on the Illumina NovaSeq X Plus platform with 100-nt single-end reads. Ribosomal RNA depletion, library construction, and raw data generation were performed by the University of Illinois Roy J. Carver Biotechnology Center Sequencing Core Facility. For cross-species test, *Xenopus* and zebrafish ribosomal RNAs were depleted together using the FastSelect HMR and Fish rRNA Removal Kit (Qiagen). Sequencing libraries were prepared from fractionated RNA using the Watchmaker Stranded mRNA library kit (Watchmaker Genomics). Libraries were then sequenced on the Illumina NovaSeq X Plus platform with 150-nt paired-end reads.

#### Fractionation m^6^A-RIP-seq library construction

Fifty zebrafish fully-grown oocytes, and embryos at the 2-cell and 256-cell stages were fractionated as described above. RNAs were extracted using TRIzol reagent. Fractionation m^6^A-RIP-seq libraries were prepared using a modified antibody-based m^6^A RNA immunoprecipitation protocol adapted from the EpiNext CUT-seq m^6^A RNA Enrichment Kit (EpigenTek) and NEBNext UltraExpress® RNA Library Prep Kit (NEB). The m⁶A IP libraries were prepared using a modified antibody-based m⁶A RNA immunoprecipitation protocol adapted from the EpiNext CUT-seq m⁶A RNA Enrichment Kit (EpiGentek). The corresponding input libraries were prepared from matched fractionated RNAs using the NEBNext UltraExpress® RNA Library Prep Kit (NEB). Briefly, fractionated RNAs isolated from soluble and insoluble fractions were incubated with 2 μg anti-m^6^A antibody and 4 μl magnetic affinity beads in 200 μl capture buffer at room temperature for 90 min with rotation to enrich m^6^A-containing RNAs. Following immunocapture, bead-bound RNAs were subjected to enzymatic cleavage by the addition of Nuclear Digestion Enhancer and Cleavage Enzyme Mix and incubated at room temperature for 5 min. Beads were subsequently washed three times with wash buffer and once with protein digestion buffer to remove nonspecifically bound RNAs.

m^6^A-enriched RNAs were released by proteinase K digestion at 55°C for 15 min and purified using magnetic bead-based RNA purification with ethanol-assisted RNA binding. Purified RNAs were eluted in 6.5 μl elution buffer and immediately subjected to first-strand cDNA synthesis using adaptor-containing reverse transcription reactions at 42°C for 50 min. Following reverse transcription, cDNA products were purified using magnetic beads for cleanup and subjected to adaptor ligation and second-strand library synthesis.

Sequencing libraries were amplified using indexed PCR amplification with KAPA HiFi polymerase. Amplified libraries were purified using 0.8 × magnetic bead size selection and evaluated using Agilent Bioanalyzer high-sensitivity assays. Libraries with fragment distributions centered at approximately 200–300 bp and lacking adaptor-dimer contamination were pooled and subjected to Illumina sequencing

#### RNA extraction, cDNA synthesis and qPCR

RNAs were extracted from the soluble and insoluble fractions using TRIzol reagent. Briefly, 800 μL of TRIzol was added to each soluble or insoluble fraction. *Egfp* RNA or *mCherry* RNA was added to the samples as a spike-in control before RNA extraction. Purified RNAs were then subjected to cDNA synthesis using random primers and reverse transcriptase according to standard protocols. Quantitative real-time PCR was performed using SYBR Green to assess mRNA levels. Ct values were acquired using the Applied Biosystems QuantStudio 3 Real-Time PCR System. The sequences of all primers used in this study are listed in Table S10.

#### PAT assay

Twenty zebrafish fully-grown oocytes or 256-cell stage embryos were collected for total RNA isolation. Fifty zebrafish fully-grown oocytes and 256-cell stage embryos were fractionated as described above to obtain soluble and insoluble RNA fractions for RNAs extraction. Purified RNAs were resuspended in RNase-free water at a final concentration of 250-1000 ng/μL. 4 μL of RNAs were ligated to an anchor RNA oligonucleotide (5’-phos-GGUCACCUUGAUCUGAAGC-NH2-3’) for 2h at 25 °C using T4 RNA ligase1 (NEB). Reverse transcription was performed using a universal anchor antisense primer (5’-GCTTCAGATCAAGGTGACCTTTTT-3’). To detect the poly(A) tail length, two rounds of PCR were performed. The first round was performed using a gene-specific outer forward primer and the anchor reverse primer. PCR products were purified using AMPure XP beads (Thermo Fisher). 1 μL of the diluted purified PCR products was used as the template for the second round of PCR with a gene-specific inner forward primer and the anchor reverse primer. PCR products were separated on a 3% agarose gel, and signal intensity was determined using the plot profile function in Image J. The sequences of all primers used in this study are listed in Table S10.

#### Analysis of fractionation RNA-seq during zebrafish OET

Raw sequencing reads were subjected to quality control using FastQC, followed by adapter and low-quality sequence trimming using Trimmomatic. Filtered reads were pseudo-aligned to the zebrafish transcriptome (GRCz11) together with synthetic spike-in RNA sequences using kallisto. Transcript abundance estimates were normalized based on the measured expression levels of gradient spike-in RNAs and subsequently converted into absolute RNA quantities expressed as attomole (amol) values.

- ***Identification of high-abundance genes:*** Total RNA abundance for each gene was calculated as the sum of soluble and insoluble fraction abundance values. High-abundance genes were defined based on the measurements of total abundance in the oocyte stage using a one-sample *t*-test, testing whether total abundance was significantly greater than 1 amol.
- ***Identification of stable genes:*** The differences in transcripts abundance were evaluated using paired statistical comparisons between developmental stages with thresholds of |log2FC| > 1 and *p* < 0.05. Transcripts with no significant changes in their abundance were retained for downstream RNA solubility analyses.
- ***RNA insolubility calculation:*** RNA insolubility was defined as the proportion of RNA abundance detected in the insoluble fraction relative to the total RNA abundance. We generated average values from four biological replicates while also preserving replicate-specific measurements for downstream statistical analyses.
- ***Analysis of insoluble RNA number and absolute amount during OET***: Insoluble RNAs were defined as transcripts with insolubility values greater than 60%. The total number and absolute amount of insoluble RNAs were calculated for each developmental stage. Differences across OET stages were assessed using one-way analysis of variance (ANOVA) followed by Tukey’s honestly significant difference (HSD) multiple-comparison test.

### Identification of dynamic RNA solubility phase-transition groups

Transcripts exhibiting dynamic changes in RNA insolubility during the OET were identified using fold-change-based criteria. “Insolubility-changing transcripts” were defined as transcripts exhibiting an absolute log2 fold change in insolubility greater than 1 between at least two developmental stages. Dynamic transcripts were clustered using k-means clustering based on their developmental insolubility trajectories to identify major RNA phase-transition patterns during OET, including S-I and I-S transitions. Transcripts without substantial insolubility changes were classified as insolubility-unchanged RNAs. Among these, transcripts with oocyte-stage insolubility values greater than 0.6 were further classified as constitutively insoluble (I-I) transcripts.

#### Reanalysis of ribosome occupancy during OET

Ribosome profiling datasets were obtained from a previously published study (Chew *et al*. 2013). Because the original analyses were performed using the Zv9/danRer7 zebrafish genome assembly, raw sequencing data were reanalyzed against the updated zebrafish reference genome (GRCz11) with minor modifications to the original computational pipeline. Briefly, raw sequencing reads were subjected to quality assessment using FastQC, followed by rRNA reads removal and read-length filtering to retain ribosome-protected fragments between 27 and 32 nucleotides. Filtered reads were aligned to the zebrafish GRCz11 reference genome using Bowtie2. Ribosome occupancy was subsequently quantified using SAMtools and normalized as reads per million mapped reads (RPM) for downstream comparative analyses across developmental stages.

Ribosome occupancy fold changes were calculated as the ratio of RPM values at the 256-cell stage relative to the 4-cell stage: 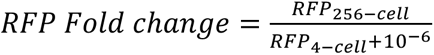. Transcripts with RFP fold change > 1.5 and detectable ribosome occupancy at the 256-cell stage were classified as exhibiting increased ribosome occupancy. Transcripts with RFP fold change < 1/1.5 and detectable ribosome occupancy at the 4-cell stage were classified as exhibiting decreased ribosome occupancy. Transcripts with fold changes between these thresholds were classified as unchanged.

#### Identification of ribosome-dependent and ribosome-independent S-I phase transition

Raw sequencing data from EDTA fractionation RNA-seq experiments were processed, quantified, and normalized using the same computational pipeline described above for fractionation RNA-seq analysis, including transcript quantification and spike-in-based normalization. Following normalization, transcripts exhibiting significant changes in soluble absolute abundance after EDTA treatment at the 256-cell stage were identified using Welch’s two-sample t-test with thresholds of *p* < 0.05 and |log2FC| > 1 and were defined as EDTA-sensitive transcripts (*n = 9,380*).

Among S-I phase transition RNAs, EDTA-insensitive transcripts (*n = 82*) were classified as Group I. Among EDTA-sensitive transcripts (*n = 1,053*), genes exhibiting decreased or unchanged ribosome occupancy were classified as Group II (*n = 465*), whereas genes showing increased ribosome occupancy were classified as Group III (*n = 184*). EDTA-sensitive transcripts that could not be detected in the ribosome profiling dataset were classified as Group IV (*n = 404*). Together, Groups I and II were considered as ribosome-independent S-I phase transition RNAs, whereas Group III was classified as the ribosome-dependent S-I phase transition group.

Using published protein expression data (Da silva pescador *et al*. 2024) and poly(A) tail length data (Chang *et al*. 2018), we calculated average protein expression levels and poly(A) tail lengths for each group. Correlations between changes in ribosome occupancy (4-cell versus 256-cell stage) and phase transition (2-cell versus 256-cell stage) of S-I RNAs were evaluated using Spearman correlation analysis.

#### Motif analysis for ribosome-dependent or -independent S-I phase transition RNAs

For ribosome-dependent and ribosome-independent S-I phase transition RNAs in zebrafish, the longest isoform of each gene was selected. The final 50 nt of the coding sequence (CDS) together with the full 3′UTR sequence were extracted for motif discovery using MEME (v5.5.9) (Bailey and elkan 1994). Motif analysis was performed using the following parameters: -mod anr - nmotifs 10 -minw 5 -maxw 10 -objfun classic -markov_order 0. Significant motifs identified from each group were further compared against the ATtRACT database (Giudice *et al*. 2016) to identify potential RBPs.

#### Analysis of cross-species fractionation RNA-seq

Raw sequencing reads were subjected to quality control using FastQC (Leggett *et al*. 2013) and adapter/quality trimming using Trimmomatic (Bolger *et al*. 2014). For cross-species quantification, zebrafish and *Xenopus* reference transcriptomes were combined into a single transcriptome index. Zebrafish cDNA sequences were based on the GRCz11 transcriptome, and *Xenopus* cDNA sequences were based on the *Xenopus laevis* v10 transcriptome. Species-specific prefixes were added to transcript identifiers before merging, with zebrafish transcripts labeled as ZF_ and *Xenopus* transcripts labeled as XL_.

Filtered paired-end reads were pseudo-aligned to the combined zebrafish and *Xenopus* transcriptome using Salmon (Patro *et al*. 2017). Reads mapping uniquely to zebrafish or *Xenopus* transcripts were retained for downstream species-specific quantification, whereas reads with multi-mapping or cross-species mapping were excluded. Transcript abundances were normalized based on gradient spike-in RNAs and converted to attomole (amol) values. Transcripts with total RNA abundance greater than 1 amol in both zebrafish and *Xenopus* datasets were retained for downstream analyses.

- ***Analysis of solubility phase transition during Xenopus oocyte maturation:*** Transcripts exhibiting increased insolubility during *Xenopus* oocyte maturation were defined as S-I phase transition RNAs if they showed log2 fold change > 1 between the oocyte and egg stages, together with *p* < 0.05 based on Student’s *t* test. Transcripts exhibiting decreased insolubility were defined as I-S phase transition RNAs if they showed log2 fold change < −1, together with *p* < 0.05.
- ***Comparison of solubility phase transitions between zebrafish and Xenopus:*** To compare solubility phase transitions between zebrafish and *Xenopus*, the same criteria used to define S-I and I-S phase transitions during *Xenopus* oocyte maturation were applied to zebrafish datasets during oocyte maturation.
- ***Identification of zebrafish and Xenopus homologs:*** Zebrafish and *Xenopus* homolog information was obtained from Xenbase (Karimi *et al*. 2018; Fortriede *et al*. 2020). Zebrafish Ensembl gene annotations were retrieved from Ensembl BioMart and merged with the Xenbase ortholog table using zebrafish Entrez Gene IDs.

#### Analysis of correlation between poly(A) tail length and insolubility of RNAs

Poly(A) tail length at 0, 1, 2, 4, 6 hpf in zebrafish were obtained from a previously study (Chang *et al*. 2018). Correlations between the insolubility and poly(A) tail length were evaluated using Spearman correlation analysis between tail-seq measurements at 0, 1, and 2 hpf and insolubility at the 2-, 32-, and 256-cell stages.

#### Analyze the correlation between m^6^A modification and solubility phase transition of RNAs

m^6^A CLIP-seq and m^6^A-seq datasets generated at 0 and 2 hpf in zebrafish were obtained from a previous study (Zhao *et al*. 2017). Published m^6^A peak information was integrated with fractionation RNA-seq datasets to evaluate the relationship between m^6^A modification and RNA solubility phase transition during the OET. Transcripts containing m^6^A peaks were classified as m^6^A-positive transcripts, whereas transcripts lacking detectable m^6^A peaks were classified as m^6^A-negative transcripts. Differences in RNA insolubility between m^6^A-positive and m^6^A-negative transcripts were evaluated using Student’s *t*-test, and the distribution of m^6^A-positive transcripts across RNA solubility transition groups was evaluated using a chi-square test.

#### Analysis of fractionation m^6^A-RIP-seq

For fractionation m^6^A-RIP-seq analysis, ribosomal RNA reads were first removed using RiboDetector. Adapter sequences and polyG tails were trimmed using fastp. Filtered reads were aligned to the zebrafish reference genome (GRCz11) using STAR, and coordinate-sorted BAM files were generated. Because the m^6^A-RIP libraries contained short inserts, only uniquely mapped reads from pulldown samples were retained for downstream peak calling to minimize ambiguous peak assignment.

m^6^A peaks were identified independently for each biological replicate and fraction sample using exomePeak2 with default parameters. Peak annotation was performed using ChIPseeker. Peaks with FDR < 0.05 located within CDS or UTR regions and reproducibly detected in both biological replicates were retained for downstream analyses. Consensus peak sets were subsequently generated across developmental stages and fractions to construct a peak-state matrix for analyzing m^6^A peak dynamics during OET.

For transcript-level classification, transcripts containing m^6^A peaks in both soluble and insoluble fractions were classified as m^6^A+ (S&I). Transcripts containing peaks exclusively in the soluble fraction were classified as m^6^A+ (S). Transcripts containing peaks exclusively in the insoluble fraction were classified as m^6^A+ (I). And transcripts without detectable m^6^A peaks were classified as m^6^A-. These transcript groups were subsequently used for comparative analyses of RNA insolubility across developmental stages.

#### Reanalysis of RNA secondary structure during the OET

RNA secondary structure datasets during zebrafish embryogenesis were obtained from a previous study (Shi *et al*. 2020). Because the original analyses were performed using the zebrafish GRCz10 transcriptome assembly, raw sequencing data were reanalyzed against the GRCz11 reference transcriptome following the original computational pipeline [https://github.com/Tsinghua-gongjing/zebrafish_structure] with minor modifications. Briefly, sequencing reads were subjected to quality control and adapter trimming, followed by additional quality filtering using Trimmomatic. The longest isoform of each gene was extracted from the GRCz11 transcriptome and used to construct a transcriptome Bowtie2 index. Filtered reads were mapped to the transcriptome using Bowtie2, followed by PCR duplicate removal and read filtering. RT-stop counts and transcript abundance were subsequently calculated, and icSHAPE reactivity scores were normalized according to the procedures described in the original study. Normalized icSHAPE scores were used for downstream RNA structure analyses in machine learning model.

#### Reanalysis of m^5^C-seq data at 2 hpf

m^5^C sequencing data at the zebrafish 2 hpf were obtained from a previous study (Liu *et al*. 2022). Because the original analyses were performed using the danRer7 genome assembly, raw sequencing data were reanalyzed against the zebrafish GRCz11 reference genome following the original computational pipeline with minor modifications. Briefly, sequencing reads were subjected to quality control and aligned to the zebrafish genome using HISAT2. To improve transcript-level mapping accuracy, unmapped reads were additionally aligned to the zebrafish transcriptome using Bowtie2, and transcriptome alignments were subsequently converted back to genomic coordinates. Genome- and transcriptome-derived BAM files were merged for downstream analyses.

Merged BAM files were used to generate nucleotide-resolution pileup files for m^5^C site calling. Candidate m^5^C sites were identified across biological replicates using the same filtering and site-calling strategy described in the original study. Reproducible m^5^C sites detected across replicates were retained for downstream analyses

### Machine learning modeling and multi-omics feature integration

- ***Construction of transcript-level multi-omics feature matrices:*** To investigate the determinants of RNA solubility phase transition during the OET, transcript-level multi-omics feature matrices were constructed by integrating sequence-derived, structural, translational, epitranscriptomic, and RBP motif-related features. For each gene, the longest isoform annotated in the zebrafish GRCz11 transcriptome was selected for downstream analyses. Transcript sequence features, including transcript length, coding sequence (CDS) length, 5′UTR length, 3′UTR length, GC content, and transcript abundance, were extracted from transcript annotations and RNA-seq datasets. Ribosome occupancy features were obtained from reanalyzed ribosome profiling datasets and normalized as reads per million mapped reads (RPM) (Chew *et al*. 2013). Poly(A) tail length measurements were integrated from previously published datasets (Chang *et al*. 2018). RNA secondary structure features were calculated from normalized icSHAPE reactivity profiles (Shi *et al*. 2020). Epitranscriptomic features, including m^6^A and m^5^C modification densities, peak numbers, and regional distributions, were integrated from m^6^A CLIP-seq (Zhao *et al*. 2017), m^6^A-seq (this work), and m^5^C (Liu *et al*. 2022) sequencing datasets. To incorporate protein-level properties of motif-associated RBPs, embryonic mRNA-bound RBPs identified by UV-crosslinking at approximately 2 hpf from a previous study (Despic *et al*. 2017) were collected. Corresponding RBP annotations, including protein abundance, predicted liquid-liquid phase separation (LLPS) propensity (Hardenberg *et al*. 2020; Hatos *et al*. 2022; Vendruscolo and fuxreiter 2022), and prion-like domain/CORE scores (Lancaster *et al*. 2014), were integrated with motif-family features. For each transcript, transcript-level RBP features were summarized based on motif-associated RBPs, including average RBP property values and burden features weighted by motif abundance.
- ***RBP motif scanning and generation of motif family feature:*** RBP-binding motif annotations were obtained from the ATtRACT database (Giudice *et al*. 2016). Motif entries from zebrafish, mouse, human, *Xenopus tropicalis*, and *Xenopus laevis* were retained, and representative motifs were selected based on motif quality scores. RBPs were assigned to motif families based on RBP identity and known binding preferences to reduce feature redundancy. Low-confidence motifs, singleton motifs, and short low-complexity poly(A)-like motifs were excluded from downstream analyses. Transcript sequences from the zebrafish GRCz11 transcriptome were scanned for exact motif matches using a custom R pipeline based on Biostrings. Briefly, RNA motif sequences were converted to DNA alphabet format, and exact motif occurrences were identified across transcript sequences using Biostrings::vmatchPattern with fixed matching. For each motif hit, transcript ID, motif identity, motif family, matched sequence, motif length, and transcript-coordinate start and end positions were recorded. Motif-family-level transcript features were subsequently generated by aggregating motif hits within each transcript. These features included motif count, regional distribution across the 5′UTR, CDS, 3′UTR, and motif-family-level positional features. The resulting motif-family features were integrated with transcript-level multi-omics features for downstream machine learning analyses.
- ***Integration of local motif context features***: To evaluate the contribution of local transcript context to RNA insolubility, we constructed a motif-context feature matrix by mapping local molecular features onto individual motif occurrences across the transcriptome. These features included local m^6^A and m^5^C modification density, and local RNA secondary structure accessibility derived from icSHAPE scores.
- ***XGBoost modeling of RNA insolubility***: Machine learning models were constructed using XGBoost regression implemented in the xgboost R package. Transcript insolubility values during the OET were used as prediction targets. Integrated transcript-level multi-omics features were used as input variables. Prior to model training, features containing missing or non-finite values were filtered or imputed as appropriate. Feature matrices were randomly partitioned into training and testing datasets. Hyperparameter optimization was performed using grid-search-based parameter tuning across multiple combinations of tree depth, learning rate, subsampling ratio, column sampling ratio, and minimum child weight. Model performance was evaluated using Pearson correlation coefficients, root mean square error (RMSE), and coefficient of determination (*R²*) between predicted and observed insolubility values in the testing dataset.
- ***SHAP interpretation and feature contribution analysis***: To interpret feature contributions to RNA insolubility prediction, SHAP values were calculated using the shapviz framework. Global SHAP importance scores were used to rank transcript-level and motif-related features contributing to RNA insolubility prediction. To further investigate regulatory features associated within Group I, II, and III, we performed SHAP analysis. Additional models were generated using combined ribosome occupancy and m^6^A states to evaluate the contribution of poly(A) tail length.
- ***Motif-context association analysis:*** To identify motif-context features associated with RNA insolubility, transcripts were ranked according to their insolubility levels and divided into high-insolubility (top quartile) and low-insolubility (bottom quartile) groups. For each motif family and context feature, the mean feature value was calculated separately for the two groups. A standardized effect size was computed as:

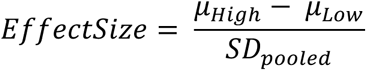

where *μ_Hi_*_)ℎ_and *μ_Lo_*_-_denote the mean feature values in the high- and low-insolubility groups, respectively, and

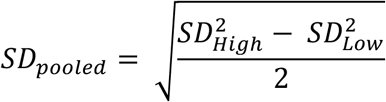

For context features whose lower values were hypothesized to promote RNA insolubility, the sign of the effect size was inverted to ensure a consistent biological interpretation, whereby positive values indicate association with increased RNA insolubility. To account for motif prevalence across transcripts, the standardized effect size was weighted by the square root of the average prevalence:

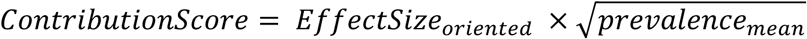

where *Prevalence_mean_* represents the average fraction of transcripts containing the corresponding motif-context feature in the two comparison groups. Statistical significance was assessed using two-sided Wilcoxon rank-sum tests comparing feature distributions between the high- and low-insolubility groups. Resulting p-values were adjusted for multiple testing using the Benjamini–Hochberg procedure, and adjusted p-values (FDR) below 0.05 were considered statistically significant.

#### GO enrichment analysis

GO enrichment analyses were performed using Metascape (Zhou *et al*. 2019). Zebrafish gene sets were analyzed using *D. rerio* as the input species, whereas *Xenopus* gene sets were converted to human orthologs and analyzed using *H. sapiens* as the input species.

#### Quantification and Statistical Analysis

All information about the statistical details is reported in the figure legends. Statistical tests and visualization for multi-omic results were performed by R studio.

## Results

### Zebrafish maternal RNAs undergo a dynamic solubility phase transition during the OET

By performing fractionation RNA-seq, we recently discovered that a subset of maternal RNAs undergoes a solubility phase transition during *Xenopus* OET (Hwang *et al*. 2023). These include RNAs encoding many RBPs and cell cycle regulators that exhibit a soluble-to-insoluble (S-I) phase transition, and germline RNAs that transition from an insoluble state to a soluble one (Hwang *et al*. 2023). To determine if the phase transition of maternal RNAs during the OET is evolutionarily conserved, we investigated zebrafish OET by fractionation RNA-seq (Figure 1A). We collected samples at multiple time points, including fully grown oocytes, ovulated (unactivated) eggs, and embryos at the 2-, 32-, and 256-cell stages (Figure 1B). To avoid complications arising from zygotically transcribed RNAs, we did not include any later time points, as ZGA occurs at the 1,000-cell stage in zebrafish. Samples were extracted in a buffer containing Triton X-100 and separated into the soluble and insoluble fractions (Figure 1A). After adding a mixture of spike RNAs into each fractionated sample, RNAs were extracted, followed by ribosomal RNA depletion, construction of random-primed libraries, and next-generation sequencing.

**Fig 1.**
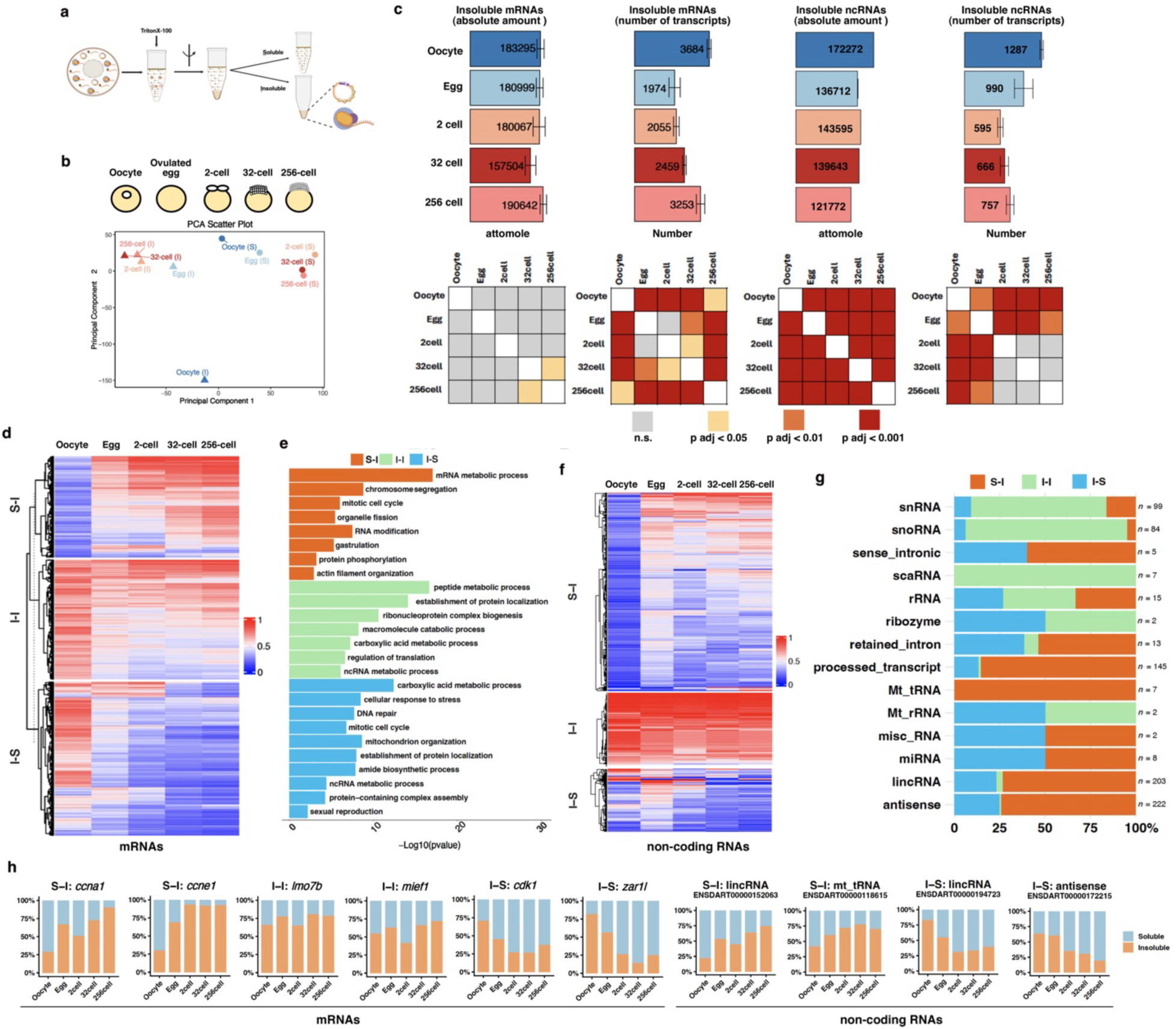
Solubility phase transition of maternal RNAs during zebrafish OET. **a.** A schematic drawing shows the procedure of fractionation RNA-seq. Created in BioRender. Chen, S. (2027) https://BioRender.com/0ni7lfu. **b**. PCA of zebrafish fractionation RNA-seq datasets across OET. PCA was performed using stage-averaged absolute RNA amount (amol) values from soluble and insoluble fractions of fully grown oocytes, unactivated eggs, and embryos at the 2-, 32-, and 256-cell stages. **c**. Quantification of insoluble mRNAs and ncRNAs during zebrafish OET. Bar graphs show the total absolute amount and number of insoluble mRNAs and ncRNAs in the oocyte, egg, and embryos at the 2-cell, 32-cell, and 256-cell stages. Insoluble transcripts were defined as RNAs with insolubility values greater than 0.6. Statistical significance was assessed using one-way ANOVA followed by Tukey’s HSD multiple-comparison test. Heatmaps indicate adjusted P values from pairwise stage comparisons. **d.** Heatmaps show the insolubility of S-I, I-I, and I-S mRNAs during OET. **e.** Bar plot from GO enrichment analysis of S-I, I-I, and I-S maternal mRNAs. Bar lengths indicate enrichment significance (−log10 adjusted p-value). **f.** Heatmaps show RNA insolubility dynamics of S-I, I-I, and I-S non-coding RNAs during zebrafish OET. **g.** Distribution of ncRNA biotypes among S-I, I-I, and I-S phase-transition groups. Numbers on the right indicate the total number of transcripts in each ncRNA category. **h.** RT-qPCR validation of mRNAs and ncRNAs exhibiting S-I, I-I, and I-S phase-transition patterns. Stacked bar plots show the proportions of RNA detected in soluble and insoluble fractions across zebrafish OET stages.

To investigate the RNA solubility phase transition, we first filtered out transcripts whose expression levels were too low (<1 attomole) or changed during the OET (Figure S1A, S1B). We found that the insoluble fraction of the oocyte was significantly different from other stages (Figure 1B, Figure S1C, and Table S1), indicating that the most dramatic solubility phase transition happens during oocyte maturation in zebrafish. Interestingly, the absolute amount of insoluble mRNAs remained largely unchanged across all five developmental stages (Figure 1C). In addition, the insolubility of the majority of transcripts remained unchanged throughout the OET (Figure S1A and S1D). To analyze the number of insoluble transcripts, we defined insoluble RNAs as those with insolubility greater than 60%. In the oocyte, we detected 3,684 insoluble mRNAs. The number of insoluble transcripts decreases to 1,974 in ovulated eggs, representing the biggest change during the OET. After fertilization, the number of insoluble transcripts gradually increases, with 2,055, 2,459, and 3,253 insoluble transcripts being detected in the 2-, 32-, and 256-cell stage embryos, respectively (Figure 1C, Table S1). Interestingly, non-coding RNAs (ncRNAs) exhibit a different pattern. The absolute amount of insoluble ncRNAs decreases gradually during the OET. The number of insoluble ncRNAs, however, decreases first, reaching the lowest number at the 2-cell stage, and stays at a low level during the cleavage (Figure 1C).

To investigate dynamic changes in the insolubility of maternal RNAs, we analyzed RNAs exhibiting dynamic changes in their insolubility by clustering (Figure S1A). In the case of mRNAs, we found 1,135 transcripts underwent an S-I phase transition, whereas 1,726 transcripts transitioned from an insoluble state to a soluble one (I-S). In addition, we detected 1,344 transcripts that remained insoluble throughout the OET (I-I) (Figure 1D, Table S1). Gene Ontology (GO) analysis revealed that mRNAs encoding RBPs, proteins important for chromosome segregation, and regulators of the mitotic cell cycle were enriched in the S-I group. I-I transcripts were involved in peptide metabolic process, establishment of protein localization, ncRNA metabolic process, etc. Consistent with our previous findings (Hwang *et al*. 2023), transcripts important for sexual reproduction exhibited I-S phase transition behavior (Figure 1E). We also identified 396 S-I, 173 I-I, and 110 I-S ncRNAs (Figure 1F, Table S1). Interestingly, most snRNAs and snoRNAs, which are assembled into snRNPs and snoRNPs with various protein complexes, fell into the I-I group. In contrast, many processed transcripts, lincRNAs, and antisense RNAs underwent an S-I phase transition. We performed fractionation RT-qPCR to validate the RNA-seq results (Figure 1H, S1E, and S1F). Indeed, *ccna1, ccne1*, lincRNA (ENSDART00000152603), mt_tRNA (ENSDART00000118615), and other candidates exhibited an S-I phase transition. The insolubility of I-I transcripts, including *imo7b, mief1*, and other candidates, remained largely stable during the OET. In contrast, *cdk1*, *zar1l*, lincRNA (ENSDART00000194723), antisense RNA (ENSDART00000172215), and other I-S transcripts were solubilized during the OET (Figure 1H, S1E, and S1F). Thus, like in *Xenopus*, zebrafish maternal RNAs undergo a solubility phase transition, indicative of dynamic interactions between maternal RNAs and protein complexes during the OET.

### The S-I phase transition of RNA during the OET can be regulated by both ribosome-dependent and ribosome-independent mechanisms

Our previous work demonstrates that remodeling of germ plasm plays an important role during the I-S phase transition of germline RNAs (Hwang *et al*. 2023). In contrast, the S-I phase transition remains poorly understood. Therefore, we chose to focus on the S-I phase transition of zebrafish maternal RNAs. In principle, the S-I phase transition of RNAs may be regulated either by increased ribosome occupancy during the OET or via the interaction between RNAs and other protein complexes. To distinguish between the ribosome-dependent and ribosome-independent RNA solubility phase transitions, we dissociated ribosomes from RNAs during fractionation using a buffer containing EDTA (Revel and hiatt 1965) (Figure 2A). The efficiency of ribosome stripping was confirmed by the redistribution of rRNAs from the insoluble fraction to the soluble fraction (Figure 2B). Moreover, fractionation RT-qPCR showed that the insolubility of S-I RNAs, including *ccna1, ccnb1*, and *hist2h3c*, was reduced under ribosome stripping conditions, whereas the I-S RNAs *dnd* and *ca15b,* two RNAs associated with the germ plasm in the embryo, were unaffected by the EDTA treatment (Figure 2C). Using this approach, we further examined the impact of ribosome-stripping on the insolubility of RNAs in 256-cell stage embryos by fractionation RNA-seq. As shown in Figure 2D, we found that the insolubility of 13,857 transcripts remained unchanged. There were 29 transcripts exhibiting increased insolubility upon EDTA treatment. In contrast, 9,380 transcripts became more soluble following the EDTA treatment (Figure 2D and Table S2).

**Fig 2.**
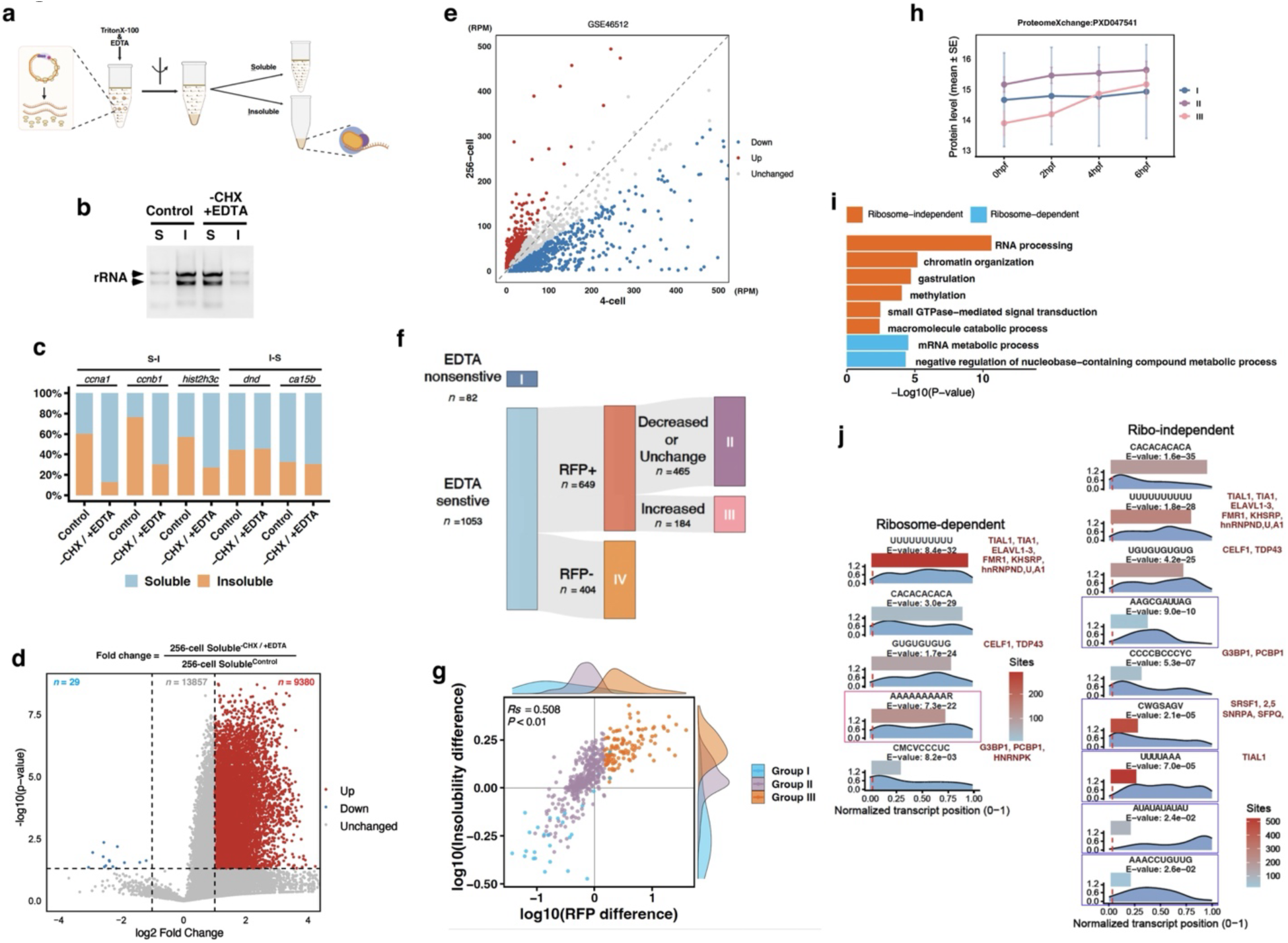
Regulation of RNA solubility phase transition by ribosome-dependent and ribosome-independent mechanisms. **a.** A schematic drawing shows the procedure of fractionation with EDTA. Created in BioRender. Chen, S. (2027) https://BioRender.com/0ni7lfu. **b**. RNA samples were separated on agarose gel to illustrate ribosomal RNA (rRNA) distribution in soluble (S) and insoluble (I) fractions following EDTA treatment. **c.** RT-qPCR shows the effects of EDTA on the insolubility of *ccna1, ccnb1, hist2h3c, dnd*, and *ca15b*. **d.** Volcano plot shows changes in the solubility of RNAs following EDTA treatment at the 256-cell stage. The amounts of soluble RNAs (amol) under control and −CHX/+EDTA conditions were compared using Welch’s two-sample t-test. Transcripts with increased, decreased, or unchanged solubility are highlighted in red, blue, and gray, respectively. **e.** Dot plot shows ribosome occupancy of transcripts at the 4-cell (x-axis) and 256-cell (y-axis) stages based on re-analysis of ribosome profiling data. Ribosome occupancy is presented as RPM. Transcripts with increased ribosome occupancy at the 256-cell stage are highlighted in red, transcripts with decreased ribosome occupancy are highlighted in blue, and unchanged transcripts are shown in gray. **f.** Sankey diagram shows the classification of S-I phase-transition RNAs into Groups I-IV according to EDTA sensitivity and ribosome occupancy during OET. **g.** Correlation between changes in ribosome occupancy and RNA insolubility among S-I phase-transition RNAs. Spearman correlation coefficient (*Rs*) and P value are shown. **h.** The abundance of proteins encoded by Groups I, II, and III transcripts during early embryogenesis based on published proteomic data (ProteomeXchange: PXD047541). Values are shown as mean ± SE. **i.** GO enrichment analysis of ribosome-independent (Groups I and II) and ribosome-dependent (Group III) S-I transcripts. Bar lengths indicate enrichment significance (−log10 adjusted P value). **j.** MEME motif enrichment analysis of ribosome-dependent and ribosome-independent S-I RNAs using the last 50 nt of the CDS and 3′UTR. Putative motif-recognizing RBPs were annotated using ATtRACT. The x-axis indicates normalized transcript position. The y-axis indicates normalized motif enrichment. Bar length represents MEME E-value. Bar color indicates the number of motif sites.

Since reduced RNA insolubility observed after EDTA treatment may be a consequence of either ribosome stripping or dissociation of other RNA-protein complexes, we combined the EDTA sensitivity data with previously reported ribosome profiling results(Chew *et al*. 2013) (Figure S2A, 2E, and Table S2) to distinguish between ribosome-dependent and ribosome-independent S-I phase transitions. We classified all S-I RNAs into four groups (Figure 2F, S2A, and Table S2). Group I comprises 82 S-I transcripts whose insolubility was not sensitive to the EDTA treatment, indicating that they were not actively translated and underwent S-I phase transitions in a ribosome-independent manner. Group II contains 465 transcripts that were sensitive to EDTA treatment. However, their ribosome occupancy either remained unchanged or even decreased from the 4-cell stage to the 256-cell stage. Because Group II transcripts exhibit an increased insolubility without a concurrent increase in ribosome occupancy, these transcripts undergo S-I phase transition via ribosome-independent mechanisms. Group III includes 184 transcripts that were both EDTA-sensitive and displayed increased ribosome occupancy during early developmental stages. Thus, their S-I phase transition is mediated by a ribosome-dependent mechanism (Figure 2F, S2B, and Table S2). In addition to these three groups, 404 transcripts were not detected in the ribosome profiling dataset. Because this may reflect the detection limits of the ribosome profiling technique, we could not conclusively determine whether the S-I phase transition of these transcripts was ribosome-dependent or ribosome-independent. Nevertheless, it is likely that many of the group IV transcripts are controlled by ribosome-independent mechanisms. These findings were further validated by independent fractionation RT-qPCR (Figure S2C). Collectively, these results demonstrate that only a subset of maternal mRNAs undergo ribosome-dependent S-I phase transition. A substantial portion of the RNA S-I phase transitions during the OET are regulated by ribosome-independent mechanisms.

We directly examined the relationship between ribosome occupancy and RNA phase transition among S-I transcripts. We found that the changes in insolubility of S-I RNAs showed a significant positive correlation with ribosome occupancy (Spearman’s *Rs* = 0.508, *p* < 0.01) (Figure 2G and Table S2). Interestingly, analysis of previously published proteomic datasets (Da silva pescador *et al*. 2024) revealed that only proteins encoded by group III transcripts showed a pronounced increase during early zebrafish embryogenesis (Figure 2H). To further characterize the biological functions associated with ribosome-independent and ribosome-dependent S-I RNAs, we performed GO analysis, combining group I and group II transcripts as ribosome-independent S-I RNAs. These transcripts were enriched for genes involved in RNA processing, chromatin organization, gastrulation, and related processes. In contrast, ribosome-dependent S-I transcripts were enriched for genes associated with mRNA metabolic processes and negative regulation of nucleobase-containing compound metabolic processes (Figure 2I). Moreover, we analyzed sequence-specific motifs in ribosome-independent (group I and II) and ribosome-dependent (group III) transcripts. We identified five motifs that were enriched in ribosome-dependent S-I transcripts. Among these, only the “AAAAAAAAAR” motif is specific to the ribosome-dependent S-I transcripts (Figure 2J, highlighted in red). The other four are also enriched in ribosome-independent S-I transcripts. Moreover, we identified 5 motifs, which were uniquely enriched in ribosome-independent S-I transcripts (Figure 2J, highlighted in blue). These motifs likely contribute to RNA S-I phase transition during the OET.

Taken together, our analysis of maternal RNA solubility phase transitions reveals highly dynamic interactions between maternal RNAs and protein complexes during the OET. While a subset of maternal RNAs becomes increasingly occupied by ribosomes and actively translated to support early embryonic development, a substantial portion of maternal RNAs appears to be recruited into unidentified protein complexes that may fine-tune gene expression during the OET.

### The capacity of maternal RNAs to engage with protein complexes changes dynamically during the OET

RNA solubility phase transition during the OET relies on precisely regulated interaction between *cis*-regulatory elements of maternal RNAs and *trans*-acting factors. Although RNA insolubility can be modulated by changes in the expression and/or post-translational modification of *trans*-acting factors during the OET (Conti and franciosi 2018; Thomalla and wolfner 2025), it remains unclear whether the capacity of maternal RNAs to engage with *trans*-acting factors also changes over time during the OET. To address this important question, we developed a cross-species assay in which zebrafish RNAs’ ability to undergo the S-I phase transition was assessed in the context of *Xenopus* oocyte maturation (Figure 3A). We reasoned that if RNAs derived from zebrafish oocytes and 256-cell stage embryos exhibit distinct insolubility in *Xenopus* oocytes and mature eggs, this would demonstrate that the ability of maternal RNAs to interact with *trans*-acting factors is dynamically regulated during the OET.

**Fig 3.**
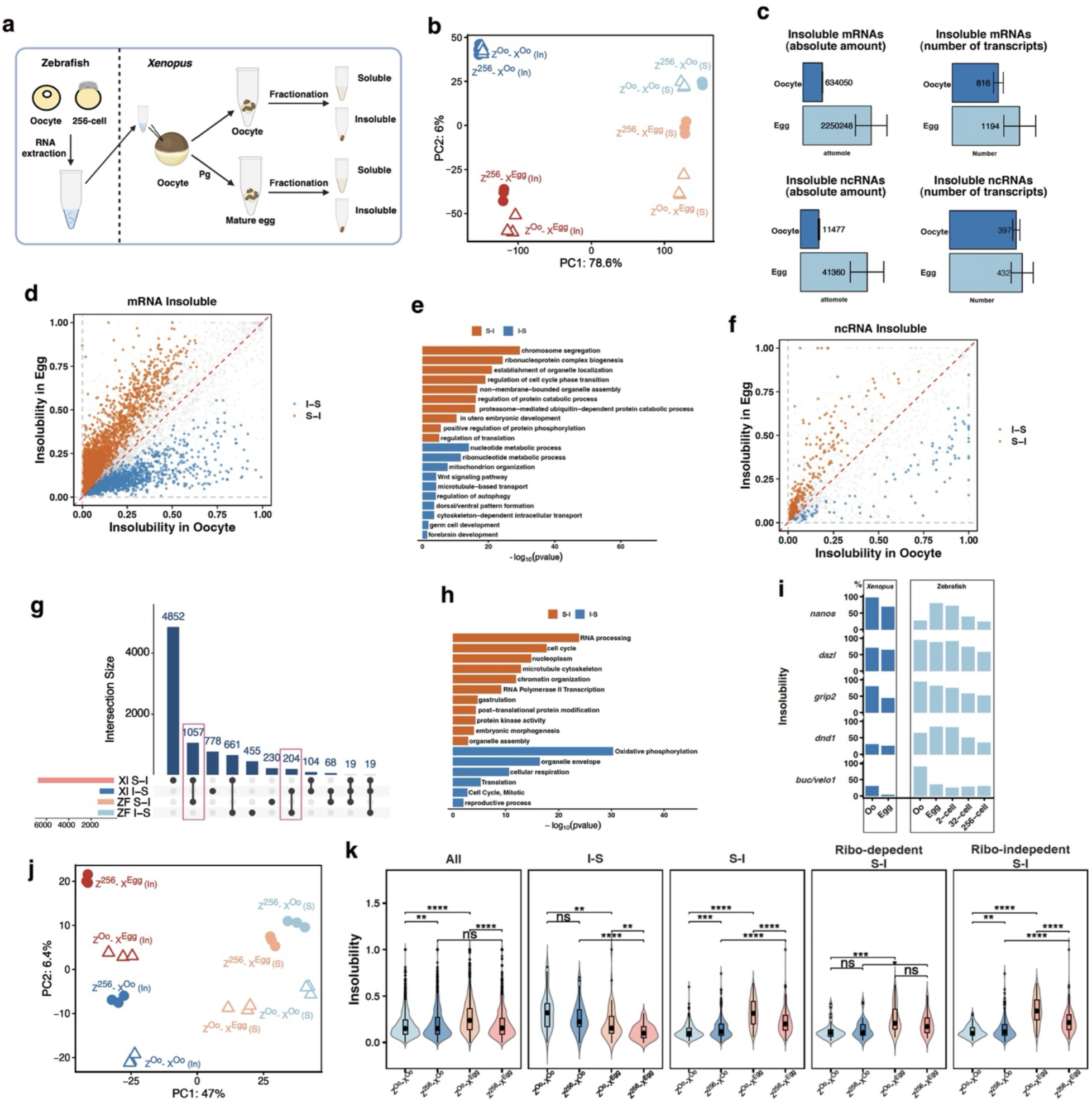
The capacity of maternal RNAs to engage with *trans*-acting factors is dynamically regulated during the OET. **a.** A schematic drawing shows the cross-species test experiments. RNAs isolated from zebrafish oocytes and 256-cell embryos were injected into *Xenopus* oocytes, followed by progesterone treatment for oocyte maturation and subsequent fractionation into soluble and insoluble RNA fractions. Created in BioRender. Chen, S. (2026) https://BioRender.com/0wo0p0q **b.** PCA of endogenous *Xenopus* transcript expression profiles reveals distinct clustering according to developmental stages and fractions. **c.** Quantification of insoluble mRNAs and ncRNAs during *Xenopus* oocyte maturation. Bar plots show the total absolute amount (amol) and number of insoluble mRNAs and ncRNAs in the oocyte and mature egg. Insoluble transcripts were defined as RNAs with insolubility values greater than 0.6. **d.** Scatter plot shows the insolubility of endogenous *Xenopus* mRNA in the oocyte and mature egg. Each dot represents a transcript. S-I RNAs – orange. I-S – blue, Unchanged transcripts – gray. **e.** GO enrichment analysis of S-I and I-S transcripts during *Xenopus* oocyte maturation. Bar lengths indicate enrichment significance (−log10 adjusted P value). **f.** Scatter plot shows ncRNA insolubility in *Xenopus* oocyte and mature egg. Each dot represents a transcript. S-I ncRNAs – orange. I-S ncRNAs – blue. Unchanged transcripts – gray. **g.** UpSet plot shows the overlap of conserved transcripts undergoing S-I and I-S phase transitions during zebrafish and *Xenopus* oocyte maturation. **h.** GO enrichment analysis of conserved transcripts undergoing S-I and I-S phase transitions in both zebrafish and *Xenopus*. The x-axis indicates enrichment significance (−log10 P value). **i.** Bar plots show that germline RNAs, including *nanos*, *dazl*, *grip2*, *dnd1*, and *buc/velo1*, undergo I-S phase transitions during *Xenopus* oocyte maturation and zebrafish OET. **j.** PCA based on the expression profiles of injected zebrafish RNAs shows distinct clustering according to developmental stages, fractions, and the developmental origins of the injected zebrafish RNAs. **k.** Violin plots comparing the insolubility of injected zebrafish RNAs derived from oocytes or 256-cell embryos during *Xenopus* oocyte maturation. Transcripts are stratified into all RNAs, I-S RNAs, S-I RNAs, ribosome-dependent S-I RNAs, and ribosome-independent S-I RNAs. Wilcoxon rank-sum test. * *p* < 0.05; ** *p* < 0.01; *** *p* < 0.001

To test the feasibility of this assay, RNAs isolated from zebrafish oocytes were injected into *Xenopus* oocytes. Half of the injected *Xenopus* oocytes were treated with progesterone (Pg) to induce oocyte maturation. The expression of *cbx2, ccnb1, hist1h2a11, ccna1,* and *tpx2*, which undergo S-I phase transition during oocyte maturation in both zebrafish (Figure S3A, upper panel) and *Xenopus* (Figure S3A, bottom panel), was analyzed by fractionation RT-qPCR. These S-I transcripts were chosen because they are abundantly expressed in zebrafish oocytes and could be reliably detected by RT-qPCR after being injected into *Xenopus* oocytes. Indeed, zebrafish *hist1h2a11, cbx2, ccnb1, tpx2*, and *hist2h3c* exhibited S-I phase transition behavior during *Xenopus* oocyte maturation (Figure S3A, middle panel), demonstrating the feasibility of studying RNA insolubility and phase transitions using this cross-species assay.

Accordingly, total RNAs isolated from zebrafish oocytes and 256-cell stage embryos were tested in the cross-species assay by fractionation RNA-seq (Figure 3A). To facilitate a more direct comparison of zebrafish RNA phase transition during *Xenopus* oocyte maturation, we adapted the zebrafish fractionation RNA-seq protocol, which uses a higher *g*-force during fractionation than the original protocol for *Xenopus* samples (Hwang *et al*. 2023; Hwang *et al*. 2024). We mapped sequencing reads to both zebrafish and *Xenopus* transcriptomes (Figure S3B). The multi-mapping rate between zebrafish and *Xenopus* was less than 0.01%, indicating successful separation of the injected zebrafish RNAs from endogenous *Xenopus* RNAs (Figure S3C). When the RNA solubility phase transition of *Xenopus* maternal RNAs during oocyte maturation was analyzed, we found that compared with our previous results (Hwang *et al*. 2023), RNAs in the current dataset showed greater enrichment in the insoluble fraction, likely due to the higher centrifugation *g*-force used during fractionation (Figure S3D). PCA result revealed that compared with soluble RNAs, transcripts in the insoluble fraction exhibited much more pronounced changes during oocyte maturation (Figure 3B and Table S3). In contrast to the pattern observed in zebrafish (Figure 1C), both the absolute amount and the number of insoluble transcripts increased during *Xenopus* oocyte maturation (Figure 3C and Table S3). Among mRNAs, 1,697 and 10,776 transcripts displayed I-S and S-I phase transition behaviors, respectively (Figure 3D and Table S3-4). GO analysis revealed that many S-I mRNAs encode proteins involved in chromosome segregation, ribonucleoprotein complex biogenesis, and related biological processes, whereas the I-S group was enriched for mRNAs encoding proteins associated with Wnt signaling, germline development, and other processes (Figure 3E). Among ncRNAs, 101 transcripts transitioned from an insoluble phase to a soluble one, whereas 302 transcripts exhibited the opposite transition pattern (Figure 3F, Table S3-4). We also assessed the evolutionary conservation of RNA solubility phase transition during oocyte maturation between *Xenopus* and zebrafish. Interestingly, we found only 1,057 and 204 homologous genes exhibited conserved S-I and I-S phase-transition behaviors, respectively (Figure 3G and Table S5). These conserved S-I RNAs encode proteins involved in RNA processing, cell-cycle regulation, post-translational protein modification, and related processes. The conserved I-S group contains RNAs encoding proteins involved in oxidative phosphorylation, cellular respiration, reproductive processes, and other functions (Figure 3H). Consistent with our previous observation (Hwang *et al*. 2023), we found that the timing of germline RNA solubilization differs between *Xenopus* and zebrafish. In *Xenopus*, germline RNAs underwent an I-S phase transition during oocyte maturation, whereas in zebrafish, germline RNA solubilization occurred during cleavage stages (Figure 3I). Thus, although RNA solubility phase transition occurs during the OET in both species, some species-specific differences nevertheless exist.

Next, we analyzed the insolubility of the injected zebrafish RNAs. Both PCA (Figure 3J) and principal coordinate analysis (PCoA) (Figure S3E, and Table S4) reveal that RNAs from zebrafish oocytes (Z^Oo^) and 256-cell embryos (Z^256^) cluster separately in *Xenopus* oocytes (-X^Oo^) and mature eggs (-X^Egg^). Notably, the difference among injected zebrafish RNAs in the insoluble fraction (In) is greater than that in the soluble fraction (S) (Figure 3J and Table S4). When the insolubility of all detected zebrafish RNAs was analyzed, small but statistically significant differences between Z^Oo^ and Z^256^ were observed in both *Xenopus* oocytes and mature eggs. While the difference between Z^Oo^-X^Oo^ and Z^256^-X^Oo^ was minimal, a more pronounced difference was detected between Z^Oo^-X^Egg^ and Z^256^-X^Egg^. Notably, Z^Oo^ and Z^256^ RNAs behaved differently during *Xenopus* oocyte maturation. The insolubility of Z^Oo^-X^Egg^ was significantly higher than that of Z^Oo^-X^Oo^. In contrast, no statistically significant difference was detected between Z^256^-X^Oo^ and Z^256^-X^Egg^ (Figure 3K, left panel, Table S5). To minimize complications arising from species-specific regulatory mechanisms, we then focused on homologous genes that exhibit conserved phase transition behavior during the OET. For I-S RNAs, the insolubilities of Z^Oo^-X^Oo^ and Z^256^-X^Oo^ were comparable in *Xenopus* oocytes. Following oocyte maturation, both samples underwent an I-S phase transition, with Z^256^-X^Egg^ displaying a more pronounced solubilization than Z^Oo^-X^Egg^ (Figure 3K, second from left). Strikingly, analysis of S-I RNAs revealed that Z^256^-X^Oo^ exhibited a higher insolubility than Z^Oo^-X^Oo^, suggesting that Z^256^ RNAs have a higher affinity for certain protein complexes in *Xenopus* oocytes compared with Z^Oo^ RNAs. Both Z^Oo^ and Z^256^ RNAs showed S-I phase transition behaviors during oocyte maturation. Interestingly, the insolubility of Z^Oo^-X^Egg^ was higher than that of Z^256^-X^Egg^ in mature eggs (Figure 3K, middle panel), demonstrating that the capacity of maternal RNAs to interact with protein complexes changes dynamically over time during the OET. We further examined the insolubilities of ribosome-dependent and ribosome-independent S-I RNAs. Ribosome-dependent Z^Oo^ and Z^256^ S-I RNAs exhibited comparable insolubility in *Xenopus* oocytes and mature eggs (Figure 3K, second from right). In contrast, analysis of ribosome-independent S-I RNAs revealed a statistically significant difference in the insolubility of Z^256^ and Z^Oo^ RNAs in *Xenopus* oocytes and mature eggs. Both ribosome-independent Z^256^ and Z^Oo^ S-I RNAs became increasingly insoluble during *Xenopus* oocyte maturation. Nevertheless, Z^Oo^ RNAs exhibited a more pronounced S-I phase transition (Figure 3K, right panel).

As transcription does not occur during the OET, the observation that subsets of RNAs from zebrafish oocytes and 256-cell stage embryos display distinct insolubility patterns in *Xenopus* oocytes and mature eggs provides strong support for the hypothesis that the capacity of maternal RNAs to interact with protein complexes is remodeled over time, which likely contributes to the RNA solubility phase transition during the OET.

### Solubility phase transition of maternal RNAs is underlined by multilayered regulatory mechanisms

The dynamic capacity of maternal RNAs to engage with protein complexes during the OET suggests that RNA–protein interactions are shaped by an evolving *cis*-regulatory architecture comprising both static sequence-encoded elements and dynamic RNA features. Previous genome-wide and machine-learning studies have demonstrated that RNA sequence contains *cis*-regulatory information that contributes to mRNA stability, including during the zebrafish maternal-to-zygotic transition (Vejnar *et al*. 2019). Building on these findings, we asked whether sequence-encoded information, together with dynamic RNA features and *trans*-acting factors, could similarly explain RNA insolubility. To systematically evaluate their individual and combinatorial contributions, we developed an XGBoost-based (Chen and guestrin 2016) machine learning framework to integrate multilayered regulatory mechanisms underlying RNA insolubility at the 2hpf (Figure 4A, Table S6-8). Notably, because many datasets are not currently available for zebrafish oocytes and earlier embryonic stages, this approach cannot yet be extended to study RNA solubility phase transition across the OET.

**Fig. 4.**
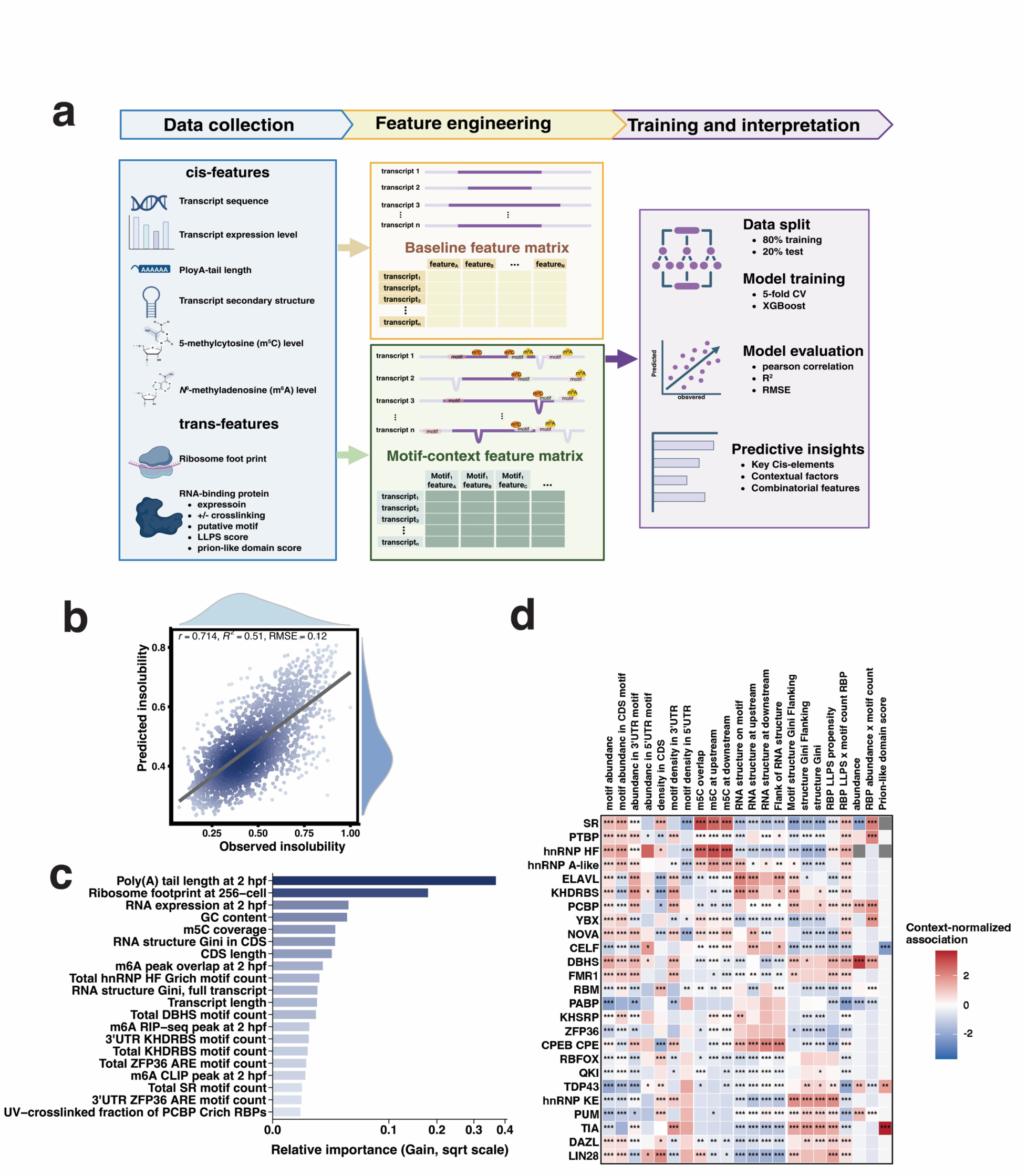
Machine learning reveals multilayered regulatory mechanisms underlying maternal RNA insolubility at 2 hpf in zebrafish embryos. **a.** Schematic overview of the machine-learning framework used to identify *cis*-elements and *trans*-regulatory features associated with RNA insolubility in 2 hpf zebrafish embryos. Created in BioRender. Chen, S. (2027) https://BioRender.com/3p02r86. **b.** Scatter plot compares observed and predicted RNA insolubility values. Model performance was evaluated using the Pearson correlation coefficient (r), the coefficient of determination (*R²*), and the root mean square error (RMSE). **c.** Feature importance derived from the XGBoost model. **d.** Heatmap shows the contribution of RNA motifs recognized by RBPs and transcript features to the insolubility of RNAs. Values represent standardized context-association scores derived from motif-context feature analyses, with red and blue indicating positive and negative contribution, respectively. Statistical significance was assessed by two-sided Wilcoxon rank-sum tests comparing transcripts with high versus low insolubility for each motif–context feature, followed by Benjamini–Hochberg correction for multiple testing. Asterisks indicate adjusted P values (FDR): * *p* < 0.05; ** *p* < 0.01; *** *p* < 0.001.

To identify the contributions of individual and combinatorial *cis*- and *trans*-regulatory features to RNA insolubility during the OET, we constructed an integrated feature matrix comprising a range of transcript-level molecular features and motif-context features (Figure 4A). *Cis*-regulatory features included RNA sequence composition, GC content, RBP-binding motifs, RNA abundance changes, and dynamic RNA properties, including secondary structure (Shi *et al*. 2020), poly(A) tail length (Chang *et al*. 2018), and epitranscriptomic modifications (Zhao *et al*. 2017; Liu *et al*. 2022). *Trans*-regulatory features included ribosome occupancy (Chew *et al*. 2013) and multiple RBP-associated properties, including RBP protein abundance (Da silva pescador *et al*. 2024), RBP-RNA interaction (Despic *et al*. 2017), liquid-liquid phase separation (LLPS) propensity (Hardenberg *et al*. 2020; Hatos *et al*. 2022; Vendruscolo and fuxreiter 2022), and prion-like domain scores (Lancaster *et al*. 2014). To systematically capture sequence-specific regulatory information, we performed transcriptome-wide motif scanning using RBP-binding motifs obtained from the ATtRACT database (Giudice *et al*. 2016) and grouped related motifs into motif families based on RBP identities and their binding sequence preferences. A total of 756 motifs representing 25 motif families were retained. Motif-context features included motif abundance, positional distribution, and local regulatory contexts, for example, whether a motif is close to specific RNA modifications or RNA structural properties. Together, this framework enabled systematic modeling of the combinatorial effects of epitranscriptomic features, sequence-specific *cis*-elements, and *trans*-acting factors on RNA insolubility.

We divided the dataset into training and independent testing sets (4:1 split stratified by RNA insolubility levels). Following hyperparameter optimization using 5-fold cross-validation on the training dataset, the final model achieved a Pearson correlation of 0.714 between the predicted and observed RNA insolubility in the independent testing dataset (Figure 4B, and Table S8), corresponding to an *R²* of 0.51. This result suggests that a substantial proportion of RNA insolubility can be explained by dynamic *cis*-regulatory features (poly(A) tail, RNA structures, and epitranscriptomics), sequence-specific *cis*-elements, and *trans*-acting factors captured by the model. The remaining unexplained variance may arise from regulatory information not included in this model, for example, transcript-specific RBP occupancy.

We next calculated the XGBoost gain scores to identify the major regulatory features associated with RNA insolubility at 2hpf (Figure 4C, Table S8). The poly(A) tail length emerged as the strongest predictor, followed by ribosome occupancy. These findings suggest that translational regulation contributes substantially to RNA insolubility. Consistent with the observations that both ribosome-dependent and ribosome-independent mechanisms contribute to S-I phase transitions of RNAs (Figure 2), the poly(A) tail length exhibited a stronger correlation with RNA insolubility as compared to ribosome occupancy. In addition to poly(A) tail length, m^6^A modification was also identified as a major contributor to RNA insolubility. The abundance of m^5^C, another RNA chemical modification, ranked among the top predictive features. However, because m^5^C frequency, i.e., the number of m^5^C sites normalized to total cytosine content, contributed minimally to model performance (Table S8), its predictive value likely reflects the GC content of the transcript rather than m^5^C modification *per se*. Notably, the Gini coefficient of RNA secondary structure within coding regions was also among the top-ranked predictive features (Figure 4C, Table S8), suggesting that transcripts with heterogeneous structural landscapes are more likely to exhibit higher insolubility. Such structural heterogeneity may facilitate selective recruitment of RBPs and promote RNA-protein interactions. Together, these analyses identify the major RNA-intrinsic features associated with RNA insolubility and suggest that tail dynamics, epitranscriptomic modifications, RNA structures, and translational regulation collectively contribute to the dynamic interaction between maternal RNAs and protein complexes.

To further investigate how sequence-specific *cis*-elements contribute to RNA insolubility, we performed the motif-context association analysis. We divided the entire transcriptome into three groups according to the insolubility of individual transcripts. We compared transcripts with high RNA insolubility (top quartile) and low RNA insolubility (bottom quartile) and calculated motif-context association scores across 25 motif families (Figure 4D, x-axis). We found that the associations of motif families to RNA insolubility were highly context-dependent (Figure 4D; Table S8). Motifs recognized by the splicing regulator-(SR-), PTBP-, hnRNPA/H/F-, ELAVL-, and KHDRBS-families of proteins were enriched in transcripts with high insolubility. These motifs are distributed throughout the transcript body. In contrast, motifs recognized by the TIA, CPEB, FMR1, ELAVL, PTBP, and PCBP protein families were preferentially enriched in the 3′UTR. These findings suggest that RNA insolubility may be influenced by the abundance and distribution of specific motifs within the entire transcript body and their density in the 3′UTR. In addition, we found that SR-, PTBP-, hnRNPA/H/F-, and ELAVL-binding motifs frequently overlapped with m^5^C sites, whereas KHDRBS-, PCBP-, DBHS-, FMR1-, hnRNPK/E-, PUM-, TIA-, and DAZL-binding motifs were associated with more complex RNA structures. This raises the intriguing possibility that the interaction between RBPs and specific transcripts may be influenced by changes in epitranscriptomic modifications and higher-order RNA structures. For *trans*-features, we found that among transcripts with high insolubility, TDP43- and PUM-binding motifs showed stronger associations with the abundance of the corresponding RBPs. When the combinatory effect of RBP abundance and the number of their binding motifs within the transcripts (RBP abundance × motif count) was considered, we found that only SR and PTBP family proteins were enriched in transcripts with high insolubility. Interestingly, while most RBPs analyzed have high liquid-liquid phase separation (LLPS) potential, only the LLPS propensities of hnRNPK/E, TDP43, PUM, TIA, and LIN28 showed a strong association with RNA insolubility. Additionally, the prion-like domain scores of TDP43 and TIA were also associated with the high insolubility of RNAs (Figure 4D).

Collectively, the above analyses demonstrate that RNA insolubility regulation is highly context-specific and is likely mediated by the interplay among multiple *cis-* and *trans-*regulatory features, including poly(A) tail dynamics, RBP-binding motifs, epitranscriptomic modifications, RNA structural landscapes, ribosome occupancy, and the biochemical properties of RBPs.

### Poly (A) tail length correlates with the insolubility and solubility phase transition of maternal RNAs during the OET

Poly (A) tail is dynamically regulated during the OET (Esencan *et al*. 2019; Ozturk 2019) and plays important roles in regulating mRNA stability and translation (Nicholson and pasquinelli 2019). Since our integrative machine-learning analysis identified poly(A) tail length as a major predictive feature of RNA insolubility at 2hpf, we next investigated how developmental changes in poly(A) tail length are associated with RNA insolubility and solubility phase transition during the OET.

Analysis of published Tail-seq datasets (Chang *et al*. 2018) (Figure 5A) revealed a strong positive correlation between poly(A) tail length and maternal RNA insolubility at 0 hpf (*R_s_*=0.53, *p* < 1.05e-212), 1 hpf (*R_s_*=0.54, *p* < 1.05e-212), and 2 hpf (*R_s_*=0.52, *p* < 1.05e-212) during zebrafish embryonic development (Figure 5B). Consistently, S-I RNAs possess significantly longer poly (A) tails than other RNAs, whereas I-S RNAs exhibit the shortest tails at 0, 1, and 2 hpf (Figure 5C). Among the S-I RNAs, group III transcripts, which are actively translated in early embryos (Figure 2F), display the shortest tails at 0 hpf, but undergo rapid tail elongation. By 4 hpf, group III transcripts have the longest poly(A) tails among all groups. Group II transcripts, which undergo an S-I phase transition in a ribosome-independent manner, also exhibit tail elongation, although to a lesser extent. In contrast, group I transcripts undergo deadenylation between 0 hpf and 2 hpf (Figure 5D). Poly(A) tail dynamics during *Xenopus* oocyte maturation have been investigated (Yang *et al*. 2020). We thus extended our analysis by determining if the poly(A) tail length correlates with the insolubility and phase transition of maternal RNAs during *Xenopus* oocyte maturation. Interestingly, we found that I-S RNAs underwent deadenylation between the MI and MII stages, whereas S-I RNAs were polyadenylated (Figure S4A). Our analysis further revealed that the insolubility of maternal RNAs poorly correlated with poly(A) tail length in fully grown oocytes (*R_s_*=0.279, *p* < 0.001) (Figure S4B, left panel). After oocyte maturation, however, a correlation between the tail length of S-I RNAs and their insolubility was detected (*R_s_*=0.413, *p* < 0.001) (Figure S4B, middle panel). Moreover, we observed a correlation between the RNA solubility phase transition and changes in poly(A) tail length (*R_s_*=0.541, *p* < 0.001). Interestingly, when the I-S RNAs were analyzed, this correlation was lost (*R_s_*=-0.152, *p* < 0.001) (Figure S4B, right panel). Thus, in both zebrafish and *Xenopus*, the poly(A) tail length correlates with both the insolubility and solubility phase transition of maternal RNAs during the OET.

**Fig 5.**
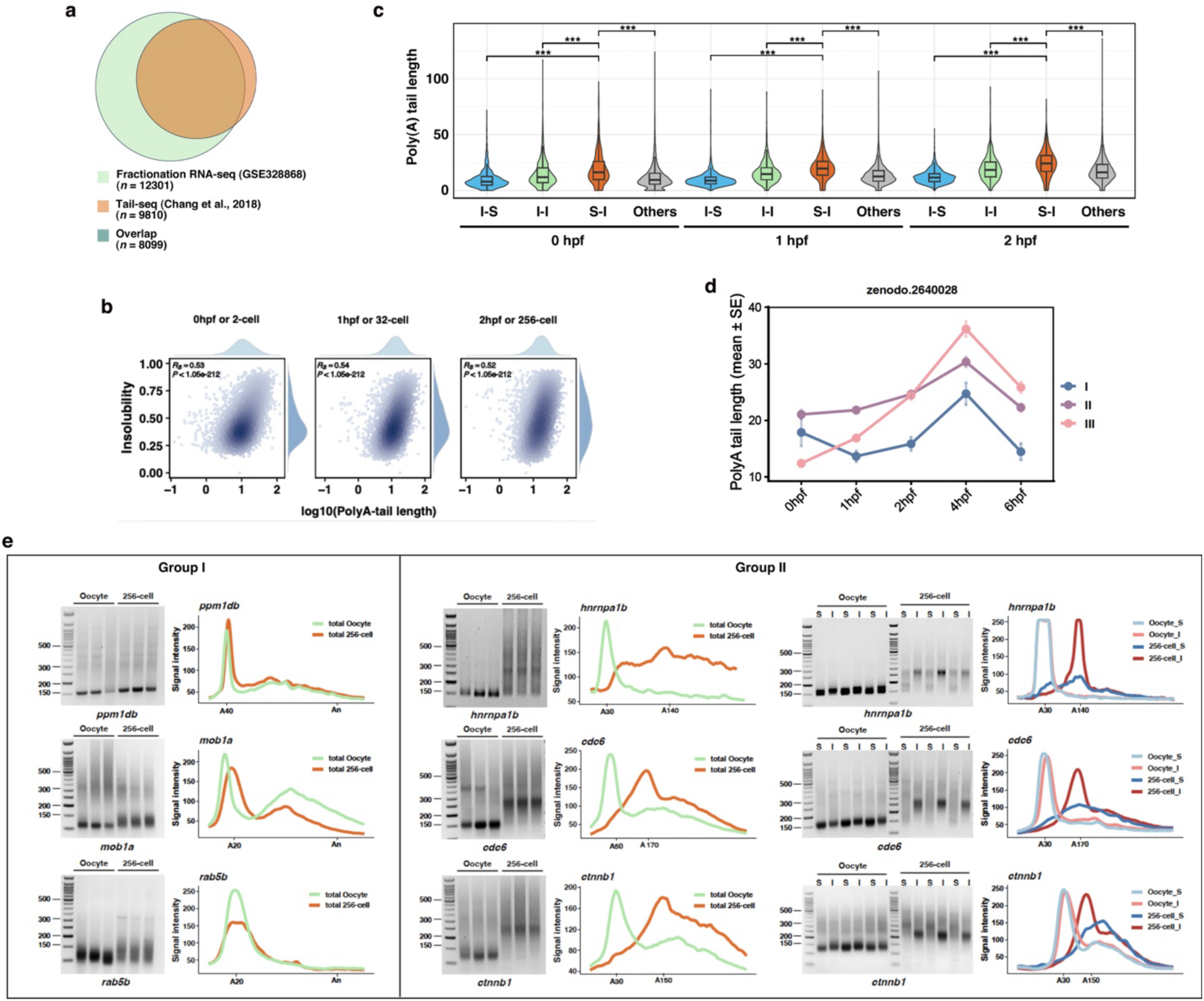
Correlation between the poly(A) tail and RNA insolubility and solubility phase transition. **a.** Venn diagram shows the overlap between transcripts detected in the zebrafish fractionation RNA-seq dataset and the published tail-seq dataset. **b.** Density scatter plots show the correlation between RNA insolubility and poly(A)-tail length. Poly(A)-tail lengths at 0, 1, and 2 hpf were compared with RNA insolubility at the corresponding 2-cell, 32-cell, and 256-cell stages, respectively. Spearman correlation coefficients and *p*-values are shown. **c.** Violin plots comparing poly(A)-tail lengths among I-S, I-I, S-I, and other transcripts at 0, 1, and 2 hpf. Statistical significance was assessed using the Wilcoxon rank-sum test. * *p* < 0.05; ** *p* < 0.01; *** *p* < 0.001. **d.** Line plots show temporal changes in poly(A) tail length of transcripts from Groups I, II, and III during zebrafish OET. Data are presented as mean ± SE. **e.** PAT assays show poly(A) tail lengths of representative Group I and Group II transcripts during zebrafish OET. Poly(A) tail lengths were compared between oocytes and 256-cell embryos. For Group II transcripts, soluble (S) and insoluble (I) fractions were analyzed separately. Representative gels and signal-intensity profiles are shown.

To further validate the relationship between poly(A) tail length and RNA insolubility identified by our transcriptome-wide analyses, we performed a poly(A) test (PAT) assay on representative transcripts from groups I and II. We selected *ppm1db*, *mob1a*, and *rab5b* as markers for group I, and *hnrnpa1b*, *cdc6*, and *ctnnb1* as markers for group II. These markers exhibit higher insolubility within their respective groups. As shown in Figure 5E, the tail length of *ppm1db*, *mob1a*, and *rab5b* remained unchanged during the OET. In contrast, *hnrnpa1b*, *cdc6*, and *ctnnb1* underwent substantial polyadenylation. Importantly, analysis of the soluble and insoluble fractions at the 256-cell stage showed that the longer, polyadenylated forms of *hnrnpa1b*, *cdc6*, and *ctnnb1* were preferentially enriched in the insoluble fraction (Figure 5E). Given that the ribosome occupancy of group II transcripts remains unchanged or decreased (Figure 2F), and that the level of their encoded proteins remains largely unchanged during cleavage (Figure 2H), the recruitment of polyadenylated group II transcripts into protein complexes likely serves as a fine-tuning mechanism to regulate their translation.

### The correlation between m^6^A modification of maternal RNAs and RNA insolubility during the OET is gradually established during the OET

Epitranscriptomic regulatory mechanisms, mediated by changes in RNA chemical modifications, are involved in post-transcriptional regulation of RNAs (Kuai *et al*. 2026). In addition to poly(A) tail length, our integrative machine-learning analysis identified RNA modifications as predictive features of RNA insolubility at 2 hpf. We therefore investigated whether dynamic epitranscriptomic regulation is associated with RNA insolubility and solubility phase transitions during the OET.

Using published RNA sequencing datasets, we investigated the correlation between RNA phase transition during the OET and epitranscriptomic mechanisms, including m^6^A modification (Zhao *et al*. 2017) and m^5^C modification (Yang *et al*. 2019). Although we failed to detect any correlation between RNA phase transition and m^5^C modification (not shown), we found that mRNAs with m^6^A modification, which regulates the decay of maternal mRNAs in vertebrate embryos (Roundtree *et al*. 2017; Zhao *et al*. 2017; Wu *et al*. 2022; Zhang *et al*. 2022b; Wang *et al*. 2023; Xiao *et al*. 2023; Zhu *et al*. 2023), exhibited higher insolubility at the 2-, 32-, and 256-cell stages (Figure 6A and B). Moreover, m^6^A-modified transcripts were more likely to show a S-I phase transition behavior during the OET (Figure 6C; Chi-square test, χ² = 265.67, *p* < 2.2 × 10⁻¹⁶), indicating that m^6^A modification correlates with the insolubility and S-I phase transition of maternal RNAs.

**Fig 6.**
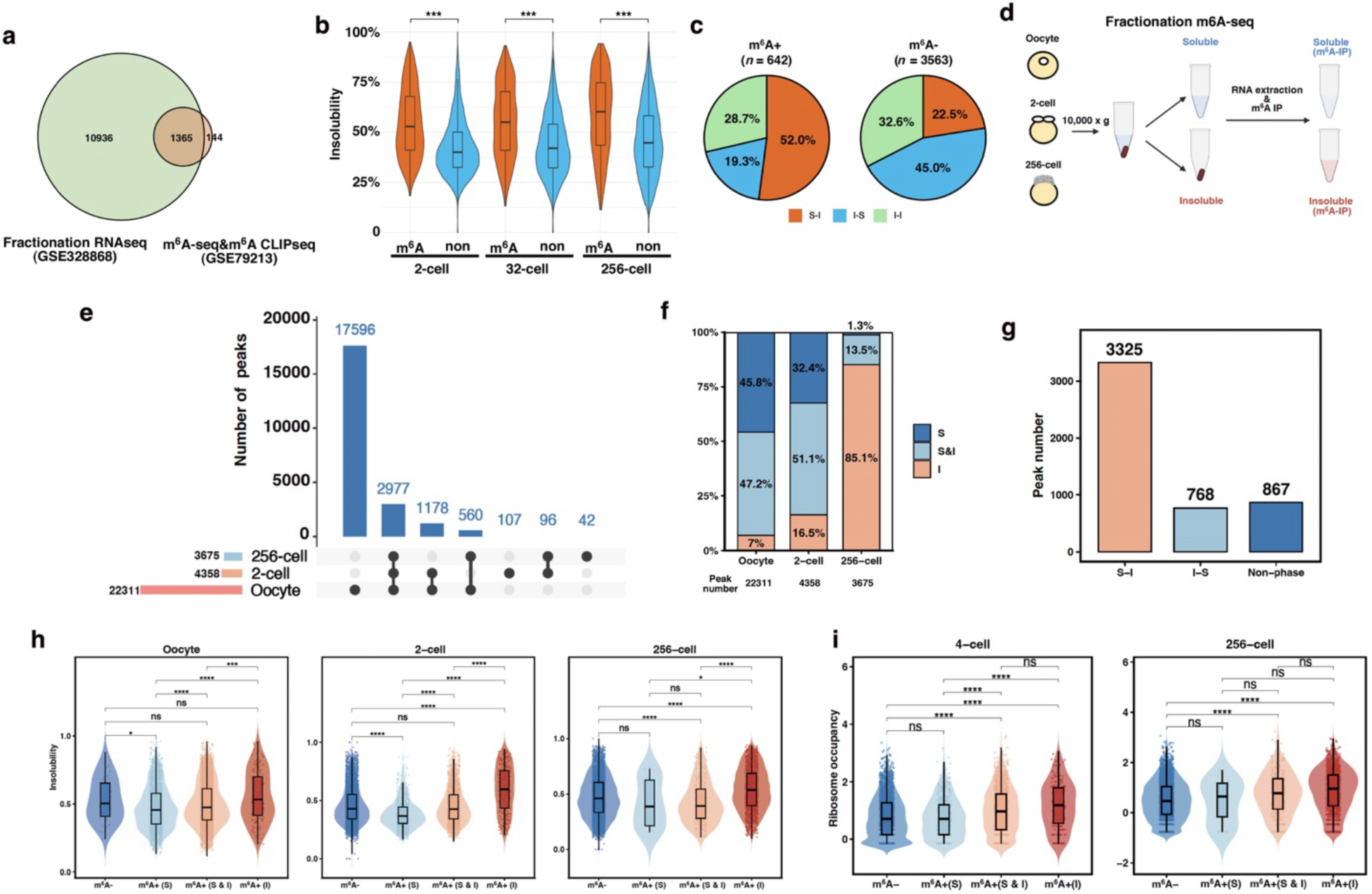
Correlation between m^6^A modification and solubility phase transition of RNAs during the OET. **a.** Venn diagram shows the overlap between transcripts detected in the fractionation RNA-seq dataset and the published m^6^A-seq & m^6^A-CLIP seq dataset. **b.** Violin plots show RNA insolubility of m^6^A-positive (m^6^A+) and m^6^A-negative (m^6^A-) transcripts at 0 hpf and 2 hpf. m^6^A status was assigned based on previously published m^6^A-seq and m^6^A-CLIP-seq datasets. **c.** Pie charts show the distribution of S-I, I-S, and I-I transcripts among m^6^A+ and m^6^A-transcripts at 0 hpf and 2 hpf. **d.** Schematic illustration of the fractionation m^6^A-RIP-seq workflow. Created in BioRender. Chen, S. (2026) https://BioRender.com/8nboxee. **e.** UpSet plot shows the overlap of m^6^A peaks identified by fractionation m^6^A-RIP-seq across the oocyte, 2-cell, and 256-cell stages. **f.** Stacked bar plots show the distribution of m^6^A peaks among soluble (S), insoluble (I), and both fractions (S&I) at the oocyte, 2-cell, and 256-cell stages. **g.** Bar plot shows the number of m^6^A peaks exhibiting S-I, I-S, and non-phase-transition behaviors identified in Figure S5B. **h** and **i.** Violin plots show RNA insolubility (**h**) and ribosome occupancy (**i**) of m^6^A-transcripts, transcripts containing m^6^A peaks in both soluble and insoluble fractions [m^6^A+(S&I)], exclusively in soluble fractions [m^6^A+(S)], and exclusively in insoluble fractions [m^6^A+(I)] during zebrafish OET. Statistical significance was assessed using the Wilcoxon rank-sum test. * *p* < 0.05; ** *p* < 0.01; *** *p* < 0.001.

To experimentally validate and further resolve the relationship between m^6^A modification and RNA insolubility during the OET, we performed a fractionation m^6^A-seq. Zebrafish oocytes and embryos at the 2- and 256-cell stages were separated into soluble and insoluble fractions, followed by m^6^A immunoprecipitation, library construction, and subsequent RNA sequencing analysis (Figure 6D and S5A). Remarkably, we found that m^6^A modification was highly dynamic during zebrafish OET. Although 22,311 m^6^A peaks were detected in oocytes, the number of m^6^A peaks decreased dramatically during development, with only 3,675 m^6^A peaks remaining by the 256-cell stage. We detected only 245 embryo-specific m^6^A peaks (Figure 6E and Table S9). Thus, while a small number of m^6^A modifications are deposited during the OET, the vast majority of m^6^A marks present in the oocyte are removed. Moreover, we found that the distribution of m^6^A peaks changed dynamically during the OET. In the oocyte, m^6^A peaks were distributed predominantly in the soluble fraction, either exclusively (45.8%) or in both the soluble and insoluble fractions (47.2%), whereas only 7% of m^6^A peaks were detected exclusively in the insoluble fraction. At the 2-cell stage, despite the extensive loss of m^6^A modifications from maternal transcripts, the overall distribution pattern remained largely unchanged, with a modest increase in the insoluble fraction-specific peaks (16.5%). By the 256-cell stage, however, the distribution shifted dramatically, with 85.1% of m^6^A peaks being localized exclusively to the insoluble fraction (Figure 6F and Table S9). To further characterize the relationship between m^6^A and RNA phase transition, we analyzed the transition of m^6^A peaks between the insoluble and soluble fractions in early embryos. Among 4,960 peaks detected in the embryos, the majority (*n* = 3,325) shifted from the soluble to the insoluble fraction. In contrast, 768 peaks exhibited an I-S phase transition. The remaining 867 peaks maintained their original state throughout early development (Figure 6G, S5B, and Table S9). These findings reveal a striking redistribution of m^6^A peaks toward the insoluble phase during the OET and suggest that m^6^A may play an important role in regulating RNA phase transitions during the OET.

To better understand how m^6^A modification influences the insolubility of maternal RNAs during the OET, we compared the insolubility of non-modified RNAs (m^6^A-) with RNAs containing m^6^A peaks being detected exclusively in the soluble fraction (m^6^A+ (S)), in both soluble and insoluble fractions (m^6^A+ (S & I)), or exclusively in the insoluble fraction (m^6^A+ (I)). Across all three developmental stages examined, the insolubility of the m^6^A+ (S) and m^6^A+ (S & I) groups was generally lower than that of the other two groups, although some pairwise comparisons did not reach statistical significance. Notably, when comparing the m^6^A- and m^6^A+ (I) groups, we found that their insolubility levels were similar in oocytes. Following fertilization, however, transcripts in the m^6^A+ (I) group became significantly more insoluble than those in the m^6^A-group (Figure 6H, S5C, and Table S9). Because RNA S-I phase transition can be mediated by a ribosome-dependent mechanism, we next examined ribosome occupancy within these four groups at the 4-cell and 256-cell stages. The m^6^A- and m^6^A+ (S) groups exhibited comparable levels of ribosome occupancy, whereas the m^6^A+ (I) and m^6^A+ (S & I) groups displayed similarly elevated ribosome occupancy. At both stages, the ribosome occupancy of the m^6^A+ (I) group was significantly higher than that of the m^6^A-group (Figure 6I, S5C, and Table S9). Taken together, the above findings suggest that m^6^A modification contributes to the regulation of RNA insolubility and S-I phase transition in a stage-specific manner during the OET, likely due to the progressive accumulation of m^6^A reader proteins, such as Rbm14 (Xiao *et al*. 2023), during early embryonic development, which allows m^6^A-modified transcripts to be selectively recruited into specific protein complexes.

## Discussion

The OET is one of the most profound developmental transitions. During the OET, a series of highly coordinated events occur sequentially, including meiotic maturation, ovulation, fertilization, and ZGA. These events are driven by maternal gene products that accumulate and are stored in fully grown oocytes, independent of transcription(Schultz *et al*. 2018). In mice (Zhao *et al*. 2020) and zebrafish (this study), more than 70% of protein-coding genes are transcribed in the fully grown oocyte. As a result, the oocyte contains a highly complex and densely packed maternal transcriptome. Precisely regulated RNA-protein interactions are thus critical for selecting specific transcripts for translation while maintaining others in a repressed state throughout the OET (Conti and franciosi 2018; Winata and korzh 2018; Sha *et al*. 2019; Ermisch and wood 2024; Lorenzo-orts and pauli 2024). Owing to technical limitations, the dynamic interactions between maternal RNAs and protein complexes during the OET have remained largely uncharacterized. Here, by investigating RNA insolubility dynamics during zebrafish OET, our work demonstrates that maternal RNAs transition between soluble and insoluble phases, revealing extensive remodeling of RNA-protein interactions throughout the OET.

RNA insolubility dynamics likely arise from differential ribosome loading, dynamic interactions between RNAs and ribonucleoprotein complexes, or remodeling of the ribonucleoprotein complexes. Indeed, we found that ribosome occupancy is a strong predictor of RNA insolubility. Intriguingly, although ribosome occupancy explains the solubility phase transition of a subset of RNAs, a substantial fraction of solubility phase transition events are driven by ribosome-independent mechanisms, revealing previously unrecognized mechanisms that fine-tune post-transcriptional regulation of maternal RNAs during the OET. We identified several *cis*-features that contribute to RNA insolubility and phase transition during the OET. These include sequence-specific *cis*-elements, which are static, and dynamic *cis*-features such as poly(A) tail dynamics and epitranscriptomics. By leveraging machine learning, we found that an intricate interplay among multilayered *cis*- and *trans*-regulatory mechanisms contributes to RNA insolubility during the OET, providing novel insights into post-transcriptional regulation of gene expression during development.

Notably, our analyses identified poly(A) tail length as the strongest predictor of RNA insolubility and phase-transition during the OET (Figure 4C, and 5B). These findings are in agreement with previous studies showing that the poly(A) tail length of maternal RNAs is tightly associated with the translational efficiency of maternal RNAs during early development (Subtelny *et al*. 2014; Xiang and bartel 2021). Interestingly, tail length emerged as a stronger predictor of RNA insolubility than ribosome occupancy (Figure 4C). This suggests that while some RNAs with long poly(A) tails are associated with ribosomes and actively translated, others may be subjected to additional layers of regulation. Indeed, unlike group III S-I transcripts, which are actively translated during cleavage, polyadenylated group II S-I transcripts lack a concurrent increase in their ribosome occupancy (Figure 2F). Group II S-I transcripts with long poly(A) tails are preferentially enriched in the insoluble fraction (Figure 5E), likely being recruited into some large ribonucleoprotein complexes or biomolecular condensates. A few group II S-I transcripts have been characterized in zebrafish. It has been reported that some of these maternal RNAs are assembled into RNA granules during the OET. Among these, *ccnb1* mRNA is assembled into repressive granules that contain the Pumilio protein. Dissociation of the *ccnb1* granule correlates with translational activation of *ccnb1* mRNA during oocyte maturation (Kotani *et al*. 2013; Horie and kotani 2016). In contrast, *pou5f3*, which is polyadenylated during cleavage, remains associated with the RNA granule throughout the OET. However, the biophysical properties of the *pou5f3* RNA granules are extensively remodeled in early embryos, providing a mechanism for fine-tuning *pou5f3* translation (Sato *et al*. 2022). In addition, germline RNAs, which belong to the I-S group, are solubilized during the OET as a consequence of germ plasm remodeling (Hwang *et al*. 2023). It seems likely that ribosome-independent RNA phase transitions constitute an important regulatory layer contributing to translational regulation during the OET.

Our results further reveal that m^6^A, a common chemical modification of RNA crucial for regulating RNA metabolism (Nachtergaele and he 2018), is another strong predictor of RNA insolubility and phase transition (Figure 4C and Figure 6). During the OET, m^6^A modification is highly dynamic and regulates both the stability and translation of maternal RNAs (Zhao *et al*. 2017; Zhang *et al*. 2022b; Wang *et al*. 2023; Li *et al*. 2025; Han *et al*. 2026; Iyyappan *et al*. 2026; Su *et al*. 2026). Consistent with these functions, our results reveal that m^6^A-modified RNAs exhibit higher insolubility and a greater tendency toward the S-I phase transition in early embryos (Figure 6B and C). Analysis of the distribution of m^6^A peaks between the soluble and insoluble phases further reveals that the majority of m^6^A peaks are removed during the OET (Figure 6E and Table S9). Despite this overall reduction, the association between m^6^A modification and RNA insolubility is established progressively during early development (Figure 6F and Table S9), likely due to the dynamic expression of m^6^A reader proteins in early embryos (Xiao *et al*. 2023). Furthermore, RNAs harboring m^6^A peaks detected in the insoluble fraction exhibit significantly higher ribosome occupancy than non-modified RNAs (Figure 6I), suggesting that m^6^A modifications promote RNA insolubility, at least in part, via a ribosome-dependent mechanism.

Interestingly, a recent study demonstrates that m^6^A can induce ribosome stalling, leading to ribosome collisions (Murakami *et al*. 2025). It will be important to determine whether a similar mechanism operates during the OET to regulate the insolubility and phase transitions of maternal RNAs. Notably, m^6^A modification may exert its effect through ribosome-independent mechanisms. The regulatory effects of m^6^A modification are mediated by m^6^A reader proteins, which can modulate the interaction of m^6^A-modified RNAs with other RBPs. In principle, when m^6^A sites overlap with or are positioned adjacent to other *cis*-regulatory elements, the binding of m^6^A readers may facilitate or interfere with the recruitment of other RBPs, thereby influencing the insolubility of m^6^A-modified RNAs. In support of this hypothesis, we found that transcripts containing SR- and hnRNP H/F-binding motifs that overlap with the m^6^A site are more likely to reside in the insoluble fraction (Figure S5D). Likewise, transcripts with Tia-, Lin28-, and hnRNP K/E-binding motifs in close proximity to heterogeneous RNA structures also exhibit higher insolubility (Figure 4D). It appears that dynamic regulatory features such as m^6^A modifications, the expression of m^6^A reader proteins, and the remodeling of RNA secondary structures cooperate to shape the complex RNA phase transition behavior during the OET.

Interestingly, although RNA phase transition occurs in both zebrafish and *Xenopus*, several species-specific differences are evident. Compared with zebrafish, *Xenopus* exhibits a substantially larger number of maternal RNAs that undergo phase transitions during oocyte maturation (Figure 3G and Table S5). Only a limited set of homologous genes displays conserved phase-transition behaviors between these two species (Figure 3G and Table S5). In addition, germline RNAs, which are markedly solubilized during *Xenopus* oocyte maturation, undergo I-S phase transition primarily during the cleavage stages in zebrafish (Figure 3I). These species differences likely reflect species-specific regulatory mechanisms. For example, the solubilization of germline RNAs is mediated by the remodeling of germ plasm matrix protein (Hwang *et al*. 2023), which occurs during *Xenopus* oocyte maturation (Hwang *et al*. 2023), but during zebrafish cleavage (Riemer *et al*. 2015). Dynamic regulatory features also appear to differ substantially among different species. For example, our data show that during zebrafish OET, the majority of m^6^A modifications are removed, whereas only a small number of new m^6^A sites are established (Figure 6E, S5E, and Table S9). A similar m^6^A dynamics occurs in bovine (Iyyappan *et al*. 2026). In contrast, during human OET, approximately 45.7% of m^6^A modifications are newly added during the OET, whereas only ∼1.3% of m^6^A modifications are removed (Figure S5E and Table S9) (Su *et al*. 2026). Mice show an intermediate pattern, with 11.8% and 13.7% of m^6^A modifications being added and removed during the OET, respectively. Thus, further work is needed to elucidate the mechanisms underlying the species-specific features of RNA phase transition during the OET.

It is worth mentioning that we investigated RNA-protein interaction dynamics during the zebrafish OET by profiling RNA solubility phase transitions using fractionation RNA-seq. Although this approach enables high-throughput analysis, its spatial and biochemical resolution is limited. Consequently, while we identified widespread and dynamic RNA phase transitions during the OET, some remodeling events may not be captured in our datasets. Furthermore, our study primarily focuses on the *cis*-regulatory mechanisms governing RNA phase transitions during the OET. Beyond ribosome occupancy, little is known about how RBPs and ribonucleoprotein complexes dynamically associate with maternal RNAs throughout this developmental transition. Future mechanistic studies will therefore be essential to elucidate the molecular basis of RNA phase transitions and the regulatory mechanisms that drive the remodeling of RNA-protein interactions during the OET. It is important to note that these OET regulatory mechanisms are likely to operate in many other biological contexts. In polarized somatic cells, such as neurons, some mRNAs are synthesized in the cell body but must be transported to the axon terminal for local translation. Dynamically regulated RNA-protein interactions play a key role in coordinating these processes. It will therefore be of great interest to determine whether the regulatory principles governing the insolubility and phase transitions of RNAs during the OET also contribute more broadly to post-translational regulation of gene expression across diverse biological settings.

Altogether, our work presented here suggests that in addition to ribosome occupancy and poly(A) tail dynamics, RNA insolubility and phase transition during the OET are governed by an intricate interplay among multiple regulatory layers, including *cis*-regulatory elements, epitranscriptomic modifications, RNA secondary structures, and RBP interactions. Collectively, our study uncovers multilayered regulatory mechanisms that shape dynamic RNA-protein interactions during the OET, providing fundamental insights into post-transcriptional gene regulation during development.

## Data and code availability

- RNA sequencing data analyzed in this study have been deposited in the Gene Expression Omnibus (GEO) with the dataset identifier GSE328868.
- Previously published datasets analyzed in this study are publicly available from the following repositories: zebrafish ribosome profiling data from GEO under accession GSE46512; m^6^A-seq and m^6^A-CLIP-seq data from GSE79213; m^5^C profiling data during zebrafish embryonic development from GSE127780; RNA secondary structure (icSHAPE) data from GSE120724; and the zebrafish embryonic mRNA-bound proteome from GSE76212. Proteomics data during zebrafish embryogenesis are available through the ProteomeXchange Consortium under accession PXD047541. Human and mouse oocyte and early embryo MeLACE-seq datasets are available under accessions HRA010484 and CRA023186, respectively. Poly(A)-tail sequencing data during zebrafish early embryonic development are available through Zenodo under DOI 10.5281/zenodo.2640028, and Poly(A)-tail sequencing data during *Xenopus* oocyte maturation are available from GEO under accession GSE134537.
- The code used in this study is publicly available through GitHub (https://github.com/atoposerwu/RNA_solubility_OET) and Zenodo (10.5281/zenodo.20932902).
- Any additional information required to reanalyze the data reported in this paper is available from the lead contact upon request.

## Acknowledgments

We thank Alvaro Hernandez and Christy Wright for ribosomal RNA depletion/library construction/RNA sequencing.

## Authors’ contribution

Study conception and design: SC, MM, BZ, and JY.

Data collection: SC, MM, DC, ML, ZW, HH, SH, HP, SC, and KP.

Analysis and interpretation of results: SC, MM, KW, WM, BZ, and JY.

Draft manuscript preparation: SC, MM, and JY.

All authors reviewed the results and approved the final version of the manuscript.

## Supplemental tables

**Table S1.** RNA insolubility across zebrafish OET, related to Figures 1C–D, 1F, S1D, and S2B

**Table S2.** Classification of EDTA-sensitive transcripts and S-I RNA subgroups in zebrafish, related to Figures 2D–G and S2B

**Table S3.** The insolubility of endogenous *Xenopus* RNAs during oocyte maturation in cross-species experiments, related to Figures 3D and 3F

**Table S4.** Changes in the insolubility of endogenous *Xenopus* RNAs and injected zebrafish RNAs during oocyte maturation, related to Figures 3D and 3F

**Table S5.** Conservation of RNA insolubility dynamics during oocyte maturation between zebrafish and *Xenopus*, related to Figure 3G and 3K.

**Table S6.** *Cis*-regulatory feature dataset at 2 hpf in zebrafish, related to Figure 4A

**Table S7.** *Trans*-regulatory feature dataset at 2 hpf in zebrafish, related to Figure 4A

**Table S8.** Feature contributions based on machine-learning models, related to Figures 4B–D **Table S9.** Zebrafish m^6^A peak annotation and dynamics during OET, related to Figures 6E–I and S5E.

**Table S10.** Primers used for RT-qPCR validation, related to Figures 1, 2, and 5

## Supplemental figure legends

**Fig S1. Characterization and validation of RNA solubility phase transitions during zebrafish OET. a.** Workflow for identification of transcripts undergoing solubility phase transitions during zebrafish OET. **b.** Heatmap shows the expression profiles of transcripts that undergo dynamic RNA solubility phase transitions during zebrafish OET. **c.** Hierarchical clustering of RNAs distributed in soluble and insoluble fractions across zebrafish OET based on transcript expression profiles. **d.** Heatmap shows the insolubility of transcripts lacking a solubility phase transition behavior during zebrafish OET. **e.** The expression levels of representative S-I, I-I, and I-S transcripts during zebrafish OET were assessed by RT-qPCR. **f.** The insolubility of representative S-I, I-I, and I-S mRNAs during zebrafish OET was assessed by fractionation RT-qPCR. **g.** The insolubility of representative S-I, I-I, and I-S ncRNAs during zebrafish OET was assessed by fractionation RT-qPCR.

**Fig S2. Classification of the S-I RNAs according to their EDTA sensitivity in fractionation and ribosome occupancy in early zebrafish embryos. a.** Workflow for the classification of ribosome-dependent and ribosome-independent RNA S-I phase transition during the OET. **b.** Heatmap shows the insolubility of Group I, II, and III transcripts during zebrafish OET. **c.** The distribution of representative Group I, II, and III transcripts in the soluble fraction was examined by fractionation RT-qPCR to validate the effects of the EDTA treatment. Statistical analysis was performed using Welch’s two-sample *t*-test. * *p* < 0.05; ** *p* < 0.01; *** *p* < 0.001.

**Fig S3. Assessing the capacity of zebrafish RNAs to interact with protein complexes during *Xenopus* oocyte maturation. a.** The insolubility of representative S-I transcripts in zebrafish (upper panel, data extracted from sequencing results) and *Xenopus* (lower panel, data extracted from sequencing results) oocyte maturation. Total RNAs isolated from zebrafish oocytes were injected into *Xenopus* oocytes. The insolubility of injected zebrafish RNAs during *Xenopus* oocyte maturation was examined by fractionation RT-qPCR (middle panel). **b.** Workflow for the cross-species test to assess the insolubility of injected zebrafish RNAs by fractionation RNA-seq. **c.** Violin plot shows the cross-species multi-mapping rates between the zebrafish and *Xenopus* transcriptomes. **d.** Venn diagrams compare S-I and I-S phase transition during *Xenopus* oocyte maturation between the current study and our previous work. **e.** Principal coordinate analysis (PCoA) of injected zebrafish transcripts and endogenous *Xenopus* transcripts. Injected zebrafish transcripts clustered significantly according to RNA fraction (PERMANOVA, *P* < 0.05), whereas endogenous *Xenopus* transcripts did not show significant clustering by RNA fraction. Betadisper analysis was not significant in all comparisons, indicating homogeneous multivariate dispersion among groups.

**Fig S4. The correlation between poly(A)-tail length and RNA solubility phase transition during *Xenopus* oocyte maturation. a.** Violin plots show poly(A)-tail length distributions of S-I, I-S, and other transcripts during *Xenopus* oocyte maturation (noPG = GV, PG90 = 90 mins after added PG, MI, Inter, and MII stages). Statistical significance was determined using the Wilcoxon rank-sum test. **b**. Density scatter plots showing the relationship between poly(A)-tail length and RNA insolubility in *Xenopus* oocytes (left) and eggs (middle). Right panel shows the correlation between the changes in poly(A)-tail length and RNA solubility phase transition during oocyte maturation. Spearman correlation coefficients (*Rs*) and *p*-values are indicated.

**Fig S5. m^6^A dynamics during zebrafish OET and cross-species comparison. a.** Workflow for processing and analyzing fractionation m^6^A-RIP-seq data. **b.** UpSet plot shows the patterns of consensus m^6^A peaks across developmental stages in the soluble and insoluble fractions. **c.** UpSet plot shows the patterns of m^6^A-modified transcripts across developmental stages in the soluble and insoluble fractions. **d.** Heatmap shows the contribution of RNA motifs and m^6^A overlap to the insolubility of RNAs. Values represent context-association scores derived from motif-context feature analyses, with red and blue indicating positive and negative contribution, respectively. **e.** Alluvial plot shows the conservation and divergence of m^6^A modifications among orthologous genes in zebrafish (GV, 2-cell, and 256-cell), bovine (GV, MII, 2-cell), mice (GV, Zygote, early 2-cell), and humans (GV, MI, MII, and Zygote).

